# NR6A1 is essential for neural crest cell specification, formation and survival

**DOI:** 10.1101/2025.01.02.630962

**Authors:** Emma L. Moore Zajic, William A. Muñoz, Jennifer F Dennis, Shachi Bhatt, Daisuke Sakai, Annita Achilleos, Ruonan Zhao, Maureen Lamb, Andrew J Price, Chris Seidel, María Tiana, Antonio Barral, Delaney Clawson, Miguel Manzanares, Paul A. Trainor

## Abstract

Neural crest cells (NCC) are a migratory progenitor cell population considered unique to vertebrates. Derived from the neuroepithelium during early embryogenesis, NCC contribute to nearly every tissue and organ system throughout the body, and disruptions in NCC development can result in congenital disorders, termed neurocristopathies. Despite decades of research, we have a poor understanding of the cellular mechanisms and signals that govern mammalian NCC formation. We discovered nuclear receptor superfamily 6 group member 1 (NR6A1/GCNF/RTR), is a novel, critical regulator of mammalian NCC specification, formation and survival. *Nr6a1* is expressed throughout the neuroepithelium in mouse embryos from E8.0 to E9.5 and briefly in newly delaminated NCC. Nr6a1 loss-of-function perturbs anterior cranial NCC formation and survival, and results in the complete agenesis of migratory NCC caudal to the first pharyngeal arch. Using mouse ESC and human iPSC differentiation into NCC, chromatin immunoprecipitation, and multiomic approaches, we demonstrate that these phenotypes are associated with perturbation of NCC specification (*Foxd3, Sox9, Sox10*) and epithelial-mesenchymal transition (EMT; *Snail1, Zeb2*), in concert with persistent expression of pluripotency-associated factors (*Oct4* and *Nanog*) in the neuroepithelium. Conditional deletion revealed that Nr6a1 is required during mid-late gastrulation, demonstrating that murine NCC specification occurs earlier than previously thought. Consistent with these observations, *in vivo* overexpression of *Oct4* in gastrulating mouse embryos disrupts NCC specification and formation. Therefore, Nr6a1 is essential for mammalian NCC formation and moreover, may function as a bimodal switch repressing pluripotency-associated factors in the neuroepithelium, while concomitantly activating NCC specifiers and regulators of EMT.

## Introduction

Neural crest cells (NCC) are a heterogeneous migratory cell population. With varying degrees of potential, NCC contribute to nearly every tissue and organ system throughout the body and are therefore essential for proper development. During early embryogenesis, NCC are specified at the neural plate border, undergo an epithelial-mesenchymal transition (EMT), delaminate from the neuroepithelium and then migrate throughout the embryo. Depending on their axial level, NCC differentiate into cartilage, bone and connective tissues in the head and neck, neurons and glia of the peripheral nervous system, smooth muscle and cardiomyocytes in the heart, endocrine cells in many organs, and pigment cells in the skin, among many others (Le Douarin and Dupin 2018). Imbued with considerable inherent plasticity, NCC have served as a conduit for intraspecies and interspecies variation, and the evolution of anatomical novelties - particularly in the head and face (Trainor and Krumlauf 2000; 2001; 2002). However, perturbations in NCC development are responsible for a diverse class of congenital anomalies known as neurocristopathies (Watt 2014, Vega-Lopez, Cerrizuela et al. 2018, Dash and Trainor 2020). Uncovering the mechanisms by which NCC regulate vertebrate development and evolution, and devising therapeutic approaches for preventing congenital neurocristopathies, requires a comprehensive understanding of the signals, switches and gene regulatory networks that govern each stage of NCC development.

Decades of research have elucidated many of the tissue interactions and signals that drive NCC specification and formation. NCC are induced at the neural plate border through contact-mediated interactions between the neural ectoderm, surface ectoderm and paraxial mesoderm (Goulding, Lumsden et al. 1993, Dickinson, Selleck et al. 1995, Liem, Tremml et al. 1995, Selleck and Bronner-Fraser 1995, Mancilla and Mayor 1996, Bonstein, Elias et al. 1998, Marchant, Linker et al. 1998, Basch and Bronner-Fraser 2006). Signals from these surrounding tissues include BMP, FGF and WNT which influence the expression of neural plate border genes *Zic1*, *Msx1* and *Pax 3/7* (Goulding, Lumsden et al. 1993, Dickinson, Selleck et al. 1995, Liem, Tremml et al. 1995, Suzuki, Ueno et al. 1997, Wilson, Graziano et al. 2000, Marchal, Luxardi et al. 2009, Patthey and Gunhaga 2014). These inductive signals together with the neural plate border factors upregulate NCC-specifier genes *Foxd3, Sox9*, and *Sox10* and EMT genes *Snai1/2*, *Zeb2* and *Twist1* to facilitate NCC specification and formation (Nieto, Sargent et al. 1994, LaBonne and Bronner-Fraser 1998, Carl, Dufton et al. 1999, LaBonne and Bronner-Fraser 2000, Dottori, Gross et al. 2001, Garcia-Castro, Marcelle et al. 2002, Spokony, Aoki et al. 2002, Aoki, Saint-Germain et al. 2003, Aybar, Nieto et al. 2003, Cheung and Briscoe 2003, Honore, Aybar et al. 2003, Nitta, Tanegashima et al. 2004, Monsoro-Burq, Wang et al. 2005, Sato, Sasai et al. 2005, Hong and Saint-Jeannet 2007, Rogers, Jayasena et al. 2012, Lander, Nasr et al. 2013, Rogers, Saxena et al. 2013). However, many of these factors and their functions, which were uncovered in non-mammalian model organisms, appear dispensable in mouse (Trainor 2005, Trainor 2005, Barriga, Trainor et al. 2015). For example, BMP4 and BMP7 function as critical inductive signals that regulate NCC specification in chicken, but when BMP4 and BMP7 are knocked out in mouse, either individually or in combination, NCC still form (Dudley, Lyons et al. 1995, Liem, Tremml et al. 1995, Luo, Hofmann et al. 1995, Winnier, Blessing et al. 1995, Selleck, Garcia-Castro et al. 1998, Solloway and Robertson 1999, Trainor, Sobieszczuk et al. 2002). Similarly, BMP2 is also not required for NCC induction in the mouse, although it is important for NCC migration (Catarina Correia et al., 2007). Neural plate border genes *Pax3* and *Pax7* are also expendable for NCC formation in mouse as is evident from single and double knockouts (Mansouri, Stoykova et al. 1996, Conway, Henderson et al. 1997, Zalc, Rattenbach et al. 2015). Finally, NCC still form, delaminate and migrate in mouse embryos even when EMT master regulators *Snai1*, *Snai2*, *Zeb2* or *Twist1* are knocked out (Van de Putte, Maruhashi et al. 2003, Murray and Gridley 2006, Van de Putte, Francis et al. 2007, Bildsoe, Loebel et al. 2009). These discrepancies between mammalian and non-mammalian species are surprising given the fundamental importance of NCC in vertebrate development, but may be attributable to developmental heterochronies, and differences in species-specific experimental approaches (Barriga, Trainor et al. 2015). Thus, the gene regulatory network underpinning mammalian NCC formation has been difficult to define and remains poorly understood.

We discovered nuclear receptor subfamily 6 group A member 1 (NR6A1) as a novel pivotal regulator of murine NCC specification, formation and survival. NR6A1 is an orphan nuclear receptor, first isolated from testes and heart tissue, that was originally named germ cell nuclear factor (GCNF) and retinoid receptor-related testis-specific receptor (RTR) (Chen, Cooney et al. 1994, Hirose, O’Brien et al. 1995). Dynamically expressed during folliculogenesis and spermatogenesis, as well as in the embryo throughout pre- and post-implantation, *Nr6a1* is critical for mouse embryo development and survival (Katz, Niederberger et al. 1997, Zhang, Akmal et al. 1998, Chung, Katz et al. 2001, Lan, Xu et al. 2009). NR6A1 loss-of-function in mouse results in an open neural tube, axial truncation, cardiac defects, and failure of chorioallantoic fusion, which leads to embryonic lethality by embryonic day (E) 10.5 (Chung, Katz et al. 2001).

We uncovered *Nr6a1* as transcriptionally downregulated in a mouse model of the craniofacial neurocristopathy, Treacher Collins syndrome (Jones et al., 2008). We then determined that *Nr6a1* is spatiotemporally expressed in the neuroepithelium and newly emigrating NCC and hypothesized that *Nr6a1* regulates NCC formation. Indeed, through loss-of-function analyses in mice we show that *Nr6a1* null embryos exhibit a deficiency in anterior cranial NCC, and complete agenesis of migrating NCC caudal to the first pharyngeal arch. These phenotypes were associated with downregulation of NCC specifiers (*Foxd3, Sox9, Sox10*) and EMT master regulators (*Snail1, Zeb2*), in concert with expansion of the neural stem cell marker Sox2, increased proliferation, and persistent expression of pluripotency-associated factors (*Oct4* and *Nanog*) in the neuroepithelium. Using mouse embryonic stem cells (ESC), and human induced pluripotent stem cells (iPSC) differentiated into NCC, as well as chromatin immunoprecipitation and multiomic approaches, we demonstrate that Nr6a1 modulates the chromatin landscape and directly binds to putative DR0 motifs in the promoter regions of NCC, EMT and pluripotency-associated factors, to regulate their expression. Furthermore, global temporal, and conditional spatiotemporal deletion revealed that Nr6a1 is specifically required during mid-late gastrulation, suggesting that NCC specification in mouse embryos commences earlier than previously recognized. Consistent with this model, *in vivo* overexpression of *Oct4* in gastrulating mouse embryos disrupts NCC specification and formation. Altogether, our work has uncovered NR6A1 as a novel regulator of mammalian NCC specification and formation. Nr6a1 may therefore function as a bimodal switch that represses pluripotency factors associated with neural stem cell maintenance and proliferation, while being required to activate a gene regulatory network of NCC specifiers and regulators of EMT.

## Results

### Spatial distribution and timing of Nr6a1 expression coincides with NCC development

Treacher Collins syndrome is a neurocristopathy caused primarily by variants in the *TCOF1* gene (Dixon, Trainor et al. 2007, Trainor, Dixon et al. 2009, Falcon, Watt et al. 2022). Characterized by downward slanting of the palpebral fissures, hypoplasia of the zygomatic complex, micrognathia and cleft palate (Dixon, Trainor et al. 2007, Trainor, Dixon et al. 2009), previous studies of mouse and zebrafish models of Treacher Collins syndrome determined these characteristic craniofacial anomalies to be caused by a deficiency in NCC. *Tcof1^+/-^*haploinsufficiency results in diminished rRNA synthesis and ribosome biogenesis, leading to increased p53-dependent apoptosis, which compromises the formation, proliferation and survival of NCC (Dixon et al., 2006; Jones et al., 2008) (Noack Watt, Achilleos et al. 2016, Sakai, Dixon et al. 2016, Watt, Neben et al. 2018, Falcon, Watt et al. 2022). Transcriptomic analyses of E8.5 *Tcof1^+/-^* and control littermate embryos revealed numerous gene expression changes in association with the molecular and cellular pathogenesis of Treacher Collins syndrome (Jones, Lynn et al. 2008). We hypothesized that downregulated genes in this mouse neurocristopathy model might also be important for normal NCC development.

Interestingly, *Nr6a1*, which was shown to regulate the transition of primitive neural stem cells to definitive neural stem cells (Akamatsu, DeVeale et al. 2009), was decreased two-fold in *Tcof1^+/-^* embryos compared to controls (Jones, Lynn et al. 2008). We therefore posited that Nr6a1 may also play a critical role in NCC development. *Nr6a1* is initially expressed in embryonic stem cells at E4.5 (Gu, LeMenuet et al. 2005) and then localizes to the embryonic ectoderm at E6.5 (Chung, Katz et al. 2001) and the neural ectoderm and primitive streak during gastrulation (E6.5 – 7.5) (Chung, Katz et al. 2001). To further characterize the spatiotemporal expression of *Nr6a1* during NCC development, we performed *in situ* hybridization on CD1 mouse embryos from E8.0 to E12.5 (Figure 1). At E8.0 *Nr6a1* continues to be expressed in the neural plate (arrowheads) and primitive streak (arrows), becoming more pronounced along the entire neuraxis from E8.5-9.5 (Figure 1A-F). Transverse histological sections of stained E8.5-E8.75 embryos showed *Nr6a1* is dynamically expressed in the dorsal most region of the neuroepithelium and in presumptive newly delaminated NCC (Figure 1E, arrows). By E10.5, *Nr6a1* is no longer expressed in the embryo, and this continues through at least E12.5 (Figure 1G-I), consistent with previous studies (Chung et al., 2001; Chang et al., 2022). Our expression analyses therefore indicated that *Nr6a1* is expressed at the right time and in the right locations to potentially function in the specification, formation and early migration of NCC.

**Figure 1.**
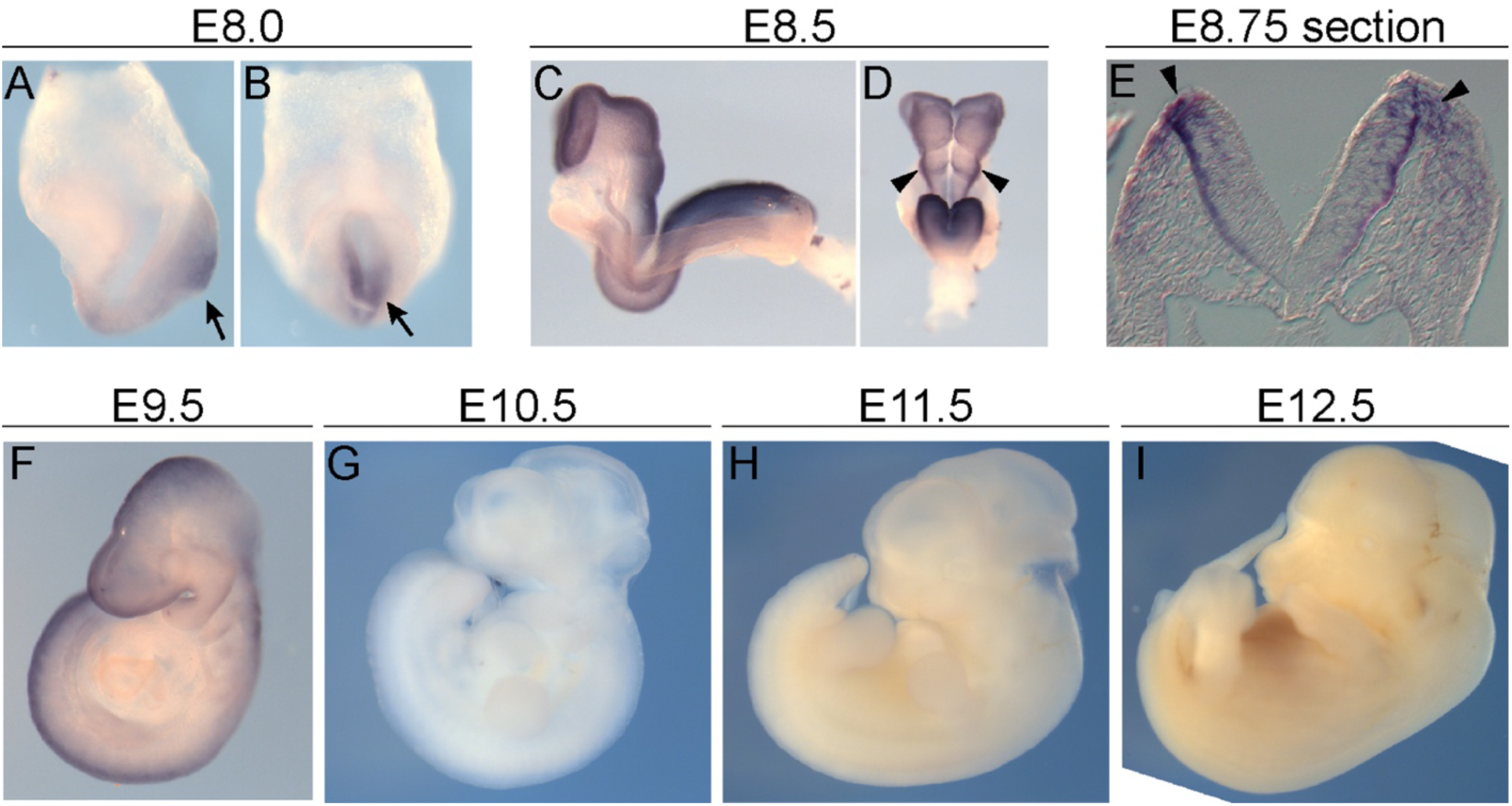
*Nr6a1* expression overlaps with regions of early NCC development. *In situ* hybridization of *Nr6a1* on CD1 embryos from E8.0-12.5. **A-B.** At E8.0, *Nr6a1* is expressed across the primitive streak and posterior regions of the embryo (arrows). **C-D.** *Nr6a1* is then expressed along the neuroepithelium and dorsal neural folds (arrowheads) at E8.5. Note that C has been reflected to maintain a similar orientation. **E.** Transverse sections across the first pharyngeal arch of E8.75 embryos shows *Nr6a1* is expressed in the dorsal region of the neuroepithelium and in cells just outside of the neuroepithelium fanning out ventrally where NCC would be located. **F.** By E9.5, *Nr6a1* expression is restricted to the neural tube. **G-I.** *Nr6a1* is no longer expressed in the embryo at E10.5 through to E12.5.

### Nr6a1 is required for NCC formation, delamination and survival

*Nr6a1* null mutant mice are embryonic lethal by E10.5 due to cardiac defects and a failure of chorioallantoic fusion (Chung, Katz et al. 2001). *Nr6a1^-/-^* embryos also exhibit significant defects in axial elongation and tail bud formation (Chung, Katz et al. 2001, Chung, Xu et al. 2006, Chang, Manent et al. 2022). To determine if *Nr6a1* is required for NCC development, we evaluated the expression of NCC-specifier genes *Foxd3*, *Sox9* and *Sox10* in E9.0 *Nr6a1^-/-^* embryos, prior to embryonic lethality (Figure 2A). *Foxd3* is one of the earliest markers of NCC specification and it maintains the potency of NCC (Mundell and Labosky 2011), while also regulating cadherin turnover to facilitate delamination by EMT (Dottori, Gross et al. 2001). Compared to wildtype littermate controls, *Foxd3* expression was absent from the NCC-derived mesenchyme in the *Nr6a1* null embryos (Figure 2A). *Sox9* is also required for NCC formation and delamination and is expressed in pre-migratory and early migratory NCC, as well as later during their chondrogenic differentiation (Ng, Wheatley et al. 1997, Zhao, Eberspaecher et al. 1997, Cheung and Briscoe 2003). We could detect only a few, if any, cells expressing *Sox9* in the craniofacial mesenchyme, with no labeled cells in the frontonasal prominences in *Nr6a1* null embryos (Figure 2A). Similarly, *Sox10*, a marker of migrating NCC (Aoki, Saint-Germain et al. 2003, Honore, Aybar et al. 2003), was absent from the pharyngeal arch mesenchyme. Only a few *Sox10* labeled cells were present in the peri-optic mesenchyme, contrasting with the widespread pattern of *Sox10*-positive NCC in the peri-optic mesenchyme and pharyngeal arch mesenchyme in controls (Figure 2A).

**Figure 2.**
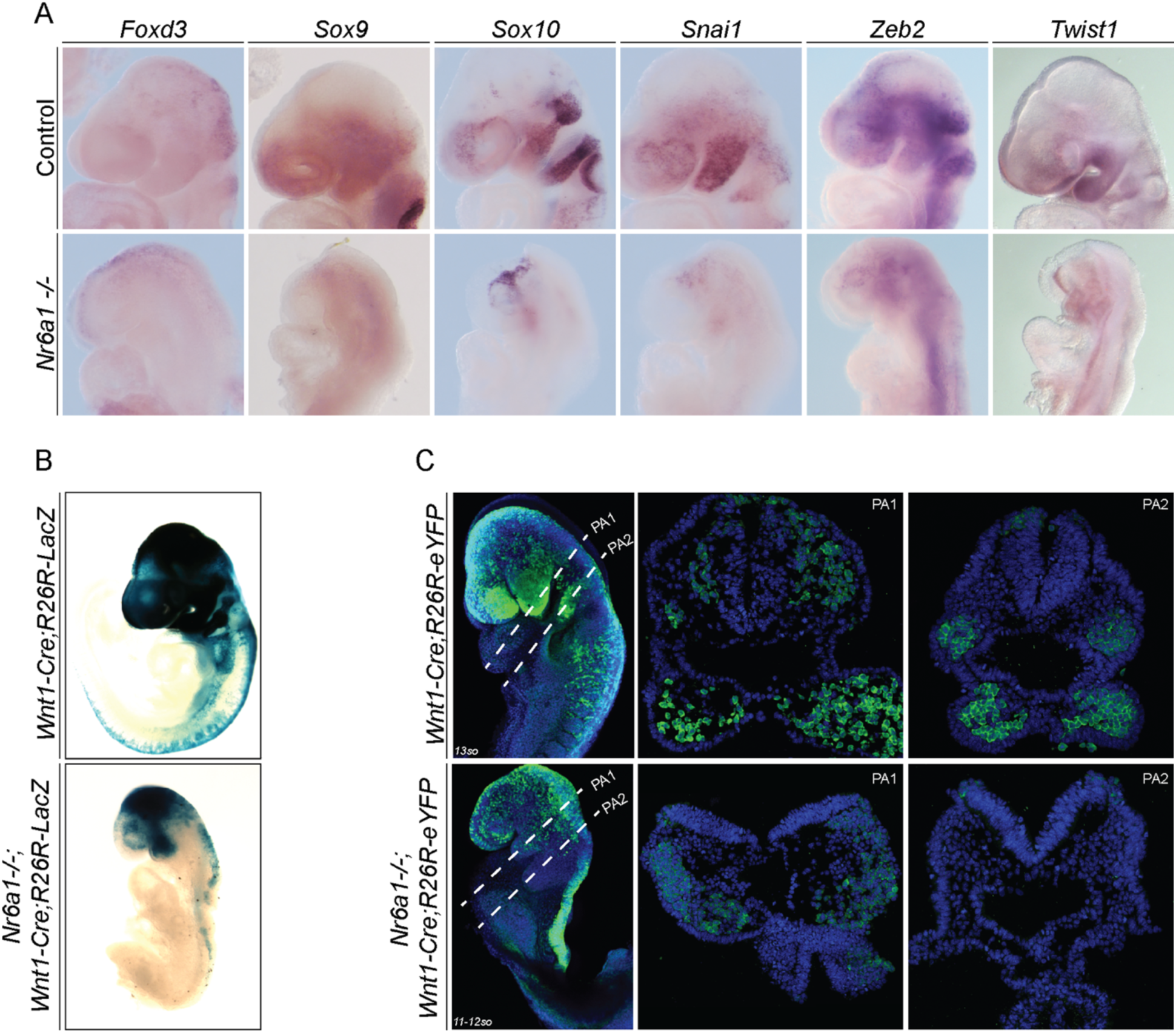
NCC-specifier and EMT genes are downregulated in the *Nr6a1* null embryos resulting in NCC deficiency. **A.** *In situ* hybridization of NCC and EMT genes on *Nr6a1* null and heterozygous littermate (control) embryos at E9.0. *Foxd3*, *Sox9*, *Sox10*, *Snai1* and *Zeb2* are downregulated in the *Nr6a1* null embryo compared to controls. *Twist1* is still expressed in the mesoderm but is not expressed in NCC until late migration. **B.** Beta-galactosidase staining of *Nr6a1^-/-^;Wnt1-Cre;R26R-LacZ* and *Wnt1-Cre;R26R-LacZ* embryos at E9.5. *Nr6a1* null embryos lack expression of LacZ caudal to the first pharyngeal arch indicating a loss of NCC. **C.** eYFP expression in *Nr6a1^-/-^;Wnt1-Cre;R26R-eYFP* and *Wnt1-Cre;R26R-eYFP* embryos at E9.0 shows a loss of NCC caudal to the first pharyngeal arch and fewer NCC in the cranial region consistent with the LacZ stained embryos in **B**. Transverse histological sections through the first pharyngeal arch (PA1) in both embryos shows a reduction in migratory NCC in the null embryos. No NCC are present in the second arch (PA2) in the null embryo.

We next evaluated whether the expression of NCC EMT master regulators *Snai1*, *Zeb2* and *Twist1* was also disrupted in *Nr6a1^-/-^*embryos. *Snai1* is typically expressed in neuroepithelial cells during NCC formation and its expression persists in delaminating and migrating NCC (Nieto, Sargent et al. 1994, Sefton, Sanchez et al. 1998, Carl, Dufton et al. 1999, Aybar, Nieto et al. 2003). However, *Snai1* expression was absent in *Nr6a1^-/-^* embryos, except for a few scattered labeled cells in the peri-optic mesenchyme (Figure 2A). *Zeb2* is also expressed in the neuroepithelium and migratory NCC (Van de Putte, Maruhashi et al. 2003, Nitta, Tanegashima et al. 2004). Although *Zeb2* was expressed in the neural plate in *Nr6a1^-/-^* embryos, it was absent from the NCC derived facial and pharyngeal mesenchyme (Figure 2A). *Twist1*, like *Snai1* and *Zeb2,* is considered a master regulator of EMT. Interestingly, *Nr6a1^-/-^* embryos exhibit *Twist1* expressing cells in the frontonasal prominences and pharyngeal arches, but to a lesser degree than in controls (Figure 2A). However, previous studies have shown that *Twist1* is not required for NCC formation but instead regulates NCC migration and differentiation during mouse embryo development (Gitelman 1997, Tavares, Izpisuja-Belmonte et al. 2001, Bildsoe, Loebel et al. 2009). Altogether, our gene expression analysis suggested *Nr6a1* is required for the expression of NCC-specifier and EMT genes during development.

Gene expression is not necessarily an indicator of lineage, but the perturbations in NCC-specifier and EMT gene expression implied that *Nr6a1* null embryos might be deficient in NCC formation and delamination. However, the expression of *Twist1* suggested there were some mesenchymal cells present in the frontonasal prominences and pharyngeal arches. Therefore, to more conclusively assess the presence or absence of NCC in *Nr6a1* null embryos, we performed lineage tracing by crossing *Wnt1-Cre* driven reporter transgenes (Chai, Jiang et al. 2000) into the background of *Nr6a1^-/-^*embryos. Interestingly, E9.0-9.5 *Nr6a1^-/-^;Wnt1-Cre;R26R-LacZ* embryos completely lacked migrating NCC caudal to the first pharyngeal arch (Figure 2B). Surprisingly, even though the expression of many NCC-specifier and EMT genes are lost in *Nr6a1^-/-^* embryos, the presence of LacZ-positive cells within the frontonasal region and first pharyngeal arch suggests some cells are still capable of delaminating and migrating from the midbrain and anterior hindbrain neuroepithelium (Figure 2B). We observed identical cranial and trunk patterns using an eYFP reporter instead of LacZ (Figure 2C). E9.0-9.5 *Nr6a1^-/-^;Wnt1-Cre;R26R-eYFP* embryos exhibited fewer migrating cranial NCC and a complete absence of migrating NCC caudal to the first pharyngeal arch. Transverse histological sections of E9.0-9.5 *Nr6a1^-/-^;Wnt1-Cre;R26R-eYFP* embryos further illustrate the diminishment of eYFP-labeled putative NCC within the first pharyngeal arch and their complete absence from the second pharyngeal arch compared to controls (Figure 2C). Given the expression of *Twist1* in *Nr6a1* null embryos, these eYFP- or LacZ-positive cells are likely to be mesenchymal (Figure 2A). However, the absence or downregulated expression of NCC-specifier and other EMT genes suggested these putative NCC were of undefined character, which raised questions about their long-term potential. TUNEL staining of E8.75 *Nr6a1^-/-^;Wnt1- Cre;R26R-eYFP* revealed visibly higher levels of cell death in *Nr6a1* mutants compared to *Wnt1-Cre;R26R-eYFP* littermate controls (Supplemental Figure 1). The cell death was predominantly located in the craniofacial mesenchyme and within regions of eYFP labeled cells (Supplemental Figure 1). Altogether, these results indicate *Nr6a1* is required for murine NCC formation and survival.

### Nr6a1 is required for the transition of neural stem cells to NCC

*Nr6a1* is predominantly expressed in the neural ectoderm during early NCC development and is required for neural tube closure (Chung, Katz et al. 2001). This raised the question of whether the deficiency in NCC formation and migration in *Nr6a1* null embryos was a secondary consequence of abnormal specification and patterning of the neural ectoderm. *Sox2* is a pan-neural marker of progenitor cells in the neural ectoderm and is sufficient to maintain their proliferation and stemness properties (Graham, Khudyakov et al. 2003, Wakamatsu, Endo et al. 2004). Through in situ hybridization, we observed that *Sox2* is expressed in the open neural plate in *Nr6a1* null embryos (Figure 3A), indicating that pan-neural specification of the neural plate occurs in *Nr6a1* mutants. *Wnt1* and *Pax3* are also expressed within the neural ectoderm, but specifically in dorsal domains that help establish dorsoventral patterning (Goulding, Lumsden et al. 1993, Dickinson, Selleck et al. 1995). Their spatiotemporal patterns of expression remain unchanged in *Nr6a1* null embryos compared to controls indicating that proper dorsal pattering is established in the neural plate of *Nr6a1* mutants. Immunostaining of transverse sections confirmed SOX2 and PAX3 were expressed in the neural ectoderm of the *Nr6a1^-/-^* embryos but also illustrated the increased thickness of the neural plate compared to controls (Figure 3B). SOX2 is a key regulator of proliferating neural progenitor cells throughout the entire anterior-posterior axis of the central nervous system from neural plate stages of embryogenesis through to adulthood (Ellis, Fagan et al. 2004). PAX3 also functions to maintain the neuroepithelium in a proliferative, undifferentiated state, allowing neurulation to proceed (Sudiwala, Palmer et al. 2019). We therefore hypothesized that the thickened neuroepithelium in the *Nr6a1* mutants was due to increased proliferation. Indeed, phospho-histone H3 (pHH3) immunostaining of transverse histological sections confirmed that *Nr6a1^-/-^*embryos exhibit considerably higher proliferation than littermate controls (Figure 3B).

**Figure 3.**
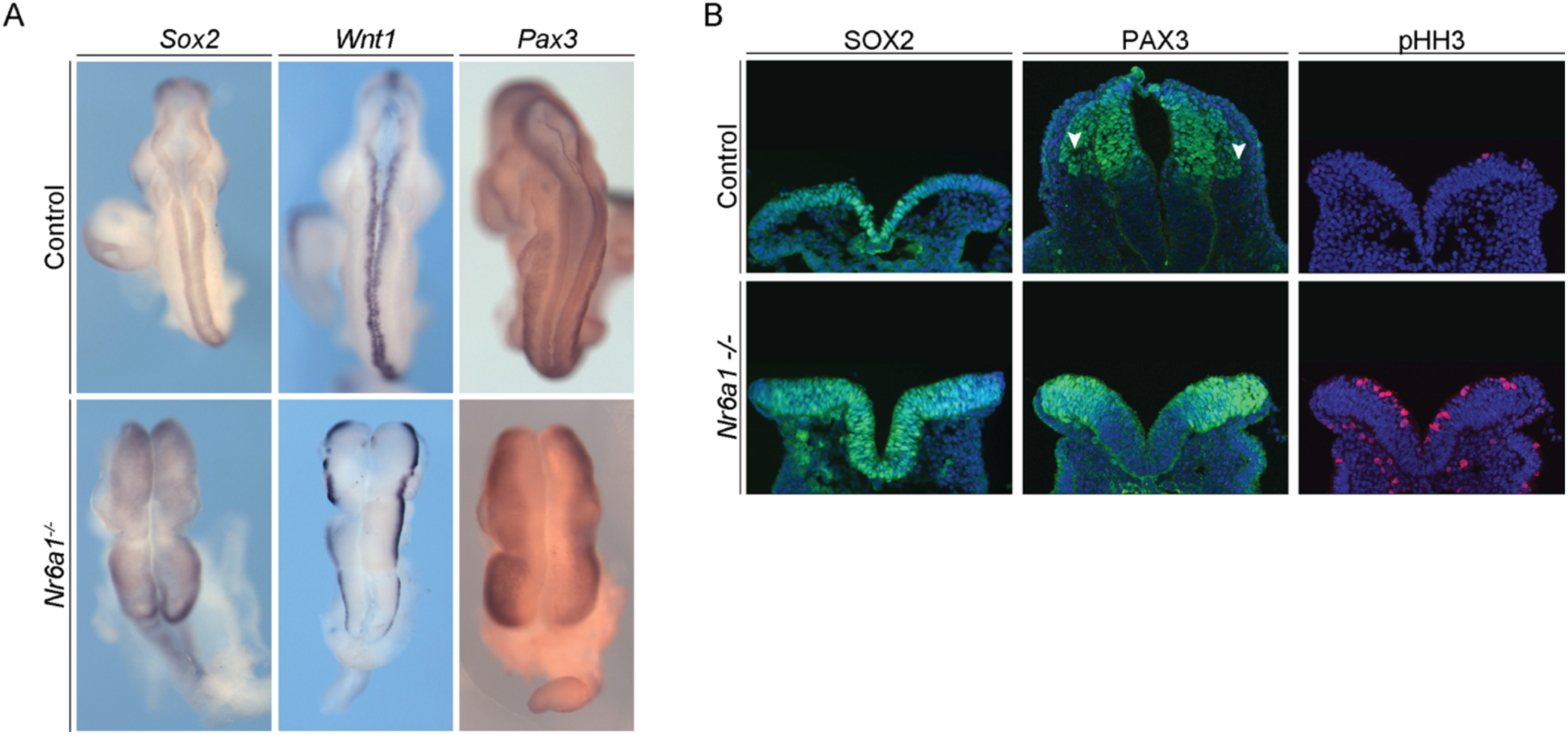
*Nr6a1* is required for the transition from neural stem to NCC. **A.** *In situ* hybridization of neuroepithelial markers in null and wildtype littermate control embryos at E9.0. *Sox2*, *Wnt1* and *Pax3* are all expressed in the *Nr6a1* null embryos, indicating the neural ectoderm has formed and patterned. **B.** Immunostaining of SOX2, PAX3 and pHH3 on transverse histological sections at E9.0. SOX2 and PAX3 are expressed in the neural plate in both null and control embryos. SOX2 expression is expanded to dorsal regions of the neural plate. PAX3 expression is maintained in the *Nr6a1* null, although no PAX3 positive migratory cells could be observed further demonstrating the NCC deficiency. More pHH3 staining can be seen in the neural plate of the *Nr6a1* null embryos suggesting the neural ectoderm is maintained in a highly proliferative state.

Additionally, we also observed that the expression of SOX2 persisted in the dorsolateral border region of *Nr6a1^-/-^* embryos compared to controls (Figure 3B). Downregulation of *Sox2* in the dorsolateral border of the neural ectoderm is essential for neural stem cells to differentiate into NCC and occurs as part of a SoxB (Sox1, Sox2) to SoxE (Sox9, Sox10) switch (Wakamatsu, Endo et al. 2004, Remboutsika, Elkouris et al. 2011). Furthermore, although the dorsal domain of PAX3 in the neural ectoderm appeared unchanged in *Nr6a1* mutants, PAX3 expression was not observed outside of the neuroepithelium, consistent with the deficiency of migratory NCC in *Nr6a1* mutants compared to littermate controls (Figure 3B).

Thus, Nr6a1 loss-of-function does not result in mis-patterning of the pan-neural or dorsal character of the neural ectoderm. Collectively, these results indicate the neural ectoderm forms and is appropriately patterned in *Nr6a1* null embryos. However, the persistence of SOX2 expression in the dorsolateral border region together with elevated proliferation, suggested that the neural ectoderm in the *Nr6a1* null mutants was maintained in a proliferative stem cell state which may restrict the neural stem cells from differentiating into NCC.

To better understand how Nr6a1 regulates NCC formation and survival, we initially performed microarray based transcriptomic analyses of E9.0 wildtype and *Nr6a1* null mutant embryos. Consistent with our in situ hybridization results (Figure 2A), the expression of NCC-specifier genes, *Foxd3*, *Sox9*, and *Sox10* was downregulated in *Nr6a1* null embryos, together with EMT master regulators *Snai1*, *Zeb2* and *Twist1* (Table 1). Although the downregulation of these factors in the microarray did not reach statistical significance, this was likely due to the entire embryo being sampled, of which the NCC are but a relatively small portion at this stage of development. Therefore, we also performed bulk RNA-sequencing, on only the anterior portion of the embryo, and again we observed considerable downregulation of NCC-specifier and EMT master regulator gene expression in *Nr6a1^-/-^* embryos compared to littermate controls (Table 1). In contrast, we found that that pluripotency associated factors such as *Oct4* and *Nanog,* were significantly upregulated in *Nr6a1^-/-^* embryos compared to littermate controls through each of the transcriptomic approaches (Table 1). Oct4 and Nanog, together with Sox2, play essential coordinated roles in preserving the pluripotency and self-renewal capacity of ESCs and adult stem cells (Nichols, Smith et al. 1998, Avilion, Nicolis et al. 2003, Wang, Rao et al. 2006). Interestingly, previous studies have shown that *Nr6a1* plays a direct role in repressing *Oct4* and *Nanog* during peri-implantation development (Fuhrmann, Chung et al. 2001, Hummelke and Cooney 2001, Gu, LeMenuet et al. 2005). To test this model in the context of NCC development, we performed a time course analysis of OCT4 expression using the *Oct4-EGFP* reporter line (Lengner, Camargo et al. 2007). Oct4-EGFP is strongly expressed and clearly observed in the epiblast at E7.5 but is then downregulated by E8.5 (Figure 4A). We then crossed *Oct4-EGFP* into the background of *Nr6a1* null embryos and observed persistent ectopic Oct4-EGFP expression throughout the neural ectoderm in E8.5 *Nr6a1^-/-^;Oct4-EGFP* embryos compared to control littermates (Figure 4B). Although GFP is an indirect readout of OCT4 activity, persistent ectopic *Oct4* expression has previously been reported in the *Nr6a1^-/-^* embryo (Fuhrmann, Chung et al. 2001). Collectively, these results indicate that OCT4 is typically downregulated in the neural ectoderm in association with NCC formation. The persistent expression of pluripotency associated genes such as *Sox2*, *Oct4* and *Nanog,* therefore suggest that the neural ectoderm in *Nr6a1* null mutants is maintained in a proliferative stem cell state which restricts neural stem cells from differentiating into NCC. Thus, *Nr6a1* may be required for the transition of neural ectoderm or neural stem cells to NCC by repressing pluripotency and stem cell maintenance genes while concomitantly being required for the expression of NCC specification and EMT genes.

**Figure 4.**
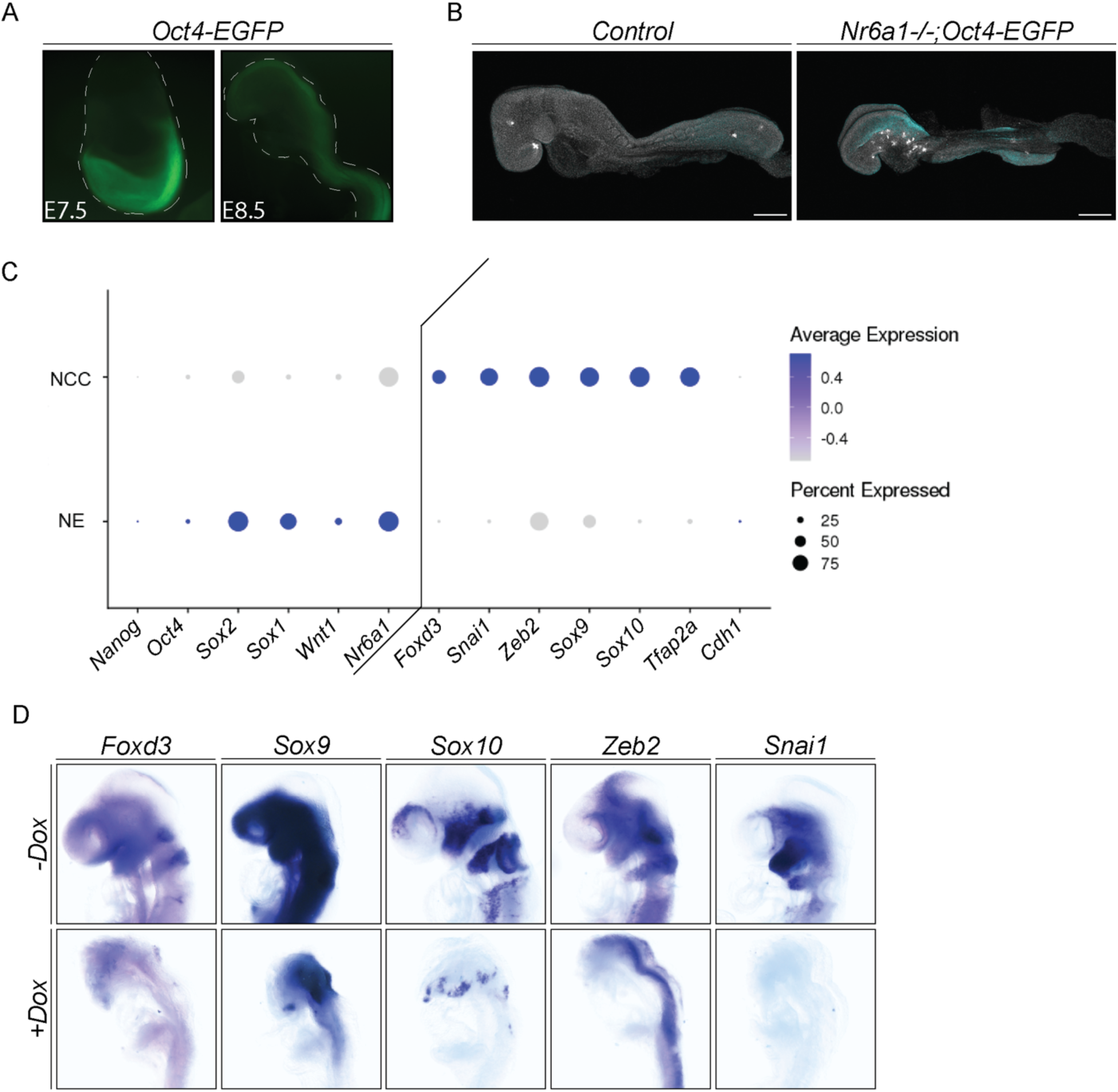
*Oct4* must be downregulated by NR6A1 prior to NCC formation. **A.** *Oct4-EGFP* expression at E7.5 and E8.5. *Oct4* is highly expressed in the embryo proper at E7.5according to high GFP expression but is downregulated by E8.5 as evidenced by the absence of GFP expression. **B.** *Oct4-EGFP* expression in *Nr6a1* null and control littermates. *Oct4* is exogenously expressed in the null compared to control embryos which do not display *Oct4* (GFP) expression. **C.** scRNA-seq of cranial tissue from *Wnt1-Cre;R26R-eYFP* and *Mef2c-F10n-LacZ* embryos shows a transcriptional signature isolating pluripotency-associated gene expression to neural ectodermal (NE) cells compared to NCC. Any expression of *Oct4* and *Nanog* is restricted to a very small population of NE cells (see dot size for percent expressed). **D.** *In situ* hybridization of NCC-specifier and EMT genes in *Oct4/rtTA* embryos treated with doxycycline (+Dox) as compared to untreated embryos (-Dox). Consistent with the *Nr6a1* null embryos, NCC and EMT marker expression is no longer present through the cranial mesenchyme of the embryos, indicating a disruption in NCC formation.

**Table 1.** Key pluripotency-associated and NCC-specifier genes are differentially expressed in *Nr6a1* null embryos compared to control littermates. scRNA-seq was performed at E8.75-9.0 on the whole embryo. Bulk RNA-seq was performed at E9.5 on the anterior of the embryos. Finally, microarray was performed at E9.0 on the whole embryo. Genes that are upregulated are in green, whereas genes that are downregulated are in red. Across all three datasets, it was found that pluripotency-associated factors *Oct4* and *Nanog* were upregulated while NCC-specifier and EMT genes were downregulated.

| scRNA-seq |  |  |  |
| --- | --- | --- | --- |
| Cell Population | Gene | log2(Nr6a1 Null/WT) | Adjusted Pvalue |
| NPB | *Oct4* | 4.107 | 5.86E-04 |
| NPB | *Nanog* | 5.379 | 3.03E-05 |
| NCC | *Sox10* | -1.961 | 7.85E-06 |

| RNA-seq |  |  |
| --- | --- | --- |
| Gene | log2(Nr6a1 Null/WT) | Adjusted Pvalue |
| *Oct4* | 5.854 | 2.54E-117 |
| *Nanog* | 4.378 | 4.70E-34 |
| *Snai1* | -0.955 | 1.71E-06 |
| *Foxd3* | -0.855 | 0.031 |
| *Sox9* | -0.293 | 0.147 |
| *Sox10* | -1.293 | 1.89E-11 |
| *Zeb2* | 0.159 | 0.484 |
| *Twist1* | -0.612 | 0.001 |

**Table 1. Key pluripotency-associated and NCC-specifier genes are differentially expressed in *Nr6a1* null embryos compared to control littermates.**
| Microarray |  |  |
| --- | --- | --- |
| Gene | log2(Nr6a1 Null/WT) | Adjusted Pvalue |
| *Oct4* | 5.379 | 0.014 |
| *Nanog* | 3.408 | 0.014 |
| *Snai1* | -0.256 | 0.373 |
| *Foxd3* | -0.355 | 0.235 |
| *Sox9* | -0.380 | 0.120 |
| *Sox10* | -0.125 | 0.574 |
| *Zeb2* | -0.325 | 0.205 |
| *Twist1* | -0.121 | 0.656 |

### NR6A1 functions by upregulating NCC-specifier and EMT genes and repressing pluripotency and stem cell maintenance

NR6A1 is an orphan member of the nuclear receptor superfamily and shares the highest homology with COUP-TF, which functions as both a transcriptional repressor and activator (Chen, Cooney et al. 1994, Hummelke and Cooney 2001). The downregulation of NCC specification and EMT genes in *Nr6a1^-/-^*embryos suggest NR6A1 could function by activating these genes during development in parallel with repressing the pluripotency-associated genes. NR6A1 binds to a double repeat of the consensus sequence AGGTCA with zero spacing (DR0) motif, to transcriptionally regulate target genes such as *Oct4* and *Nanog* (Chen, Cooney et al. 1994, Cooney, Hummelke et al. 1998, Chung, Xu et al. 2006). We therefore searched for this putative binding site or motif in NCC-specification and EMT genes to determine the potential for NR6A1 to directly regulate these genes. *Sox9*, *Sox10*, *Snai1* and *Zeb2* each possess DR0 binding sites in their promoter regions, which is consistent with downregulation of their expression in *Nr6a1* null embryos.

To further understand how NR6A1 regulates NCC-specifier and EMT genes within the neural ectoderm during NCC development, we performed single cell multiomic analyses of E8.75 *Nr6a1* null mutant and control littermate embryos. Combining single cell RNA-sequencing (scRNA-seq) with single nucleus transposase-accessible chromatin sequencing (snATAC-seq) allows for the assessment of differential gene expression and chromatin accessibility between cell populations in the *Nr6a1* null and control littermate embryos. The scRNA-seq results from the control were used to annotate cell populations (Supplemental Figure 2A-B). The neural plate border cells were identified based on high differential expression of *Wnt1*, *Pax3/7* and *Zic1 (Sauka-Spengler and Bronner-Fraser 2008, Simoes-Costa and Bronner 2015, Zhao, Moore et al. 2024)* (Supplemental Figure 2A-B). NCC were annotated based on high differential expression of the NCC-specifier genes *Foxd3*, *Sox9*, *Sox10*, *Crabp1*, *Snai1* and *Zeb2 (Trainor 2005, Stuhlmiller and Garcia-Castro 2012, Prasad, Charney et al. 2019, Zhao, Moore et al. 2024)* (Supplemental Figure 2A-B). The neural ectoderm was identified by high differential expression of *Sox2*, *Pax2*, *Otx2* and *Sox3 (Simoes-Costa and Bronner 2015, Martik and Bronner 2017, Zhao, Moore et al. 2024)* (Supplemental Figure 2A-B). Finally, neural progenitors were annotated based on the high expression of *Olig2*, *Sox2* and *Pax6* (Ericson, Rashbass et al. 1997, Graham, Khudyakov et al. 2003, Lee, Lee et al. 2005, Zhao, Moore et al. 2024) (Supplemental Figure 2A-B). Comparison of the *Nr6a1^-/-^* embryo scRNA-seq dataset to the control littermate scRNA-seq dataset revealed the NCC population was substantially decreased, reaffirming the results obtained from *Nr6a1*^-/-^;Wnt1-Cre;R26R-eYFP lineage tracing (Supplemental Figure 2C). Furthermore, the few cells that clustered as NCC in the mutant embryos exhibited a significant decrease in *Sox10* expression, while in contrast the neural plate border (NPB) cells exhibited increased *Oct4* and *Nanog* expression compared to control littermates, consistent with our previous microarray, bulk RNA-seq, in situ hybridization, immunostaining and transgenic reporter mouse analyses (Table 1).

Next, we evaluated changes in chromatin accessibility between *Nr6a1^-/-^* mutant embryos and control littermates (Supplemental Figure 3). It is well established that NR6A1 directly binds to and represses *Oct4*. Therefore, we first evaluated the chromatin accessibility of *Oct4* in the NCC and ectodermal populations in *Nr6a1^-/-^*mutant embryos compared to control littermates (Figure 5A). *Nr6a1^-/-^* embryo NCC exhibited increased chromatin accessibility compared to controls, consistent with NR6A1 loss-of-function preventing the repression of *Oct4* (Figure 5A). Furthermore, peaks illustrating the increased accessibility of open chromatin upstream of *Oct4,* aligned with the locations of putative DR0 bindings sites, demonstrating mechanistically how Nr6a1 represses *Oct4* expression (Figure 5A). We next evaluated chromatin accessibility around *Sox10*, which is significantly decreased in *Nr6a1^-/-^* embryos (Figure 5B). If NR6A1 regulates the transcription of *Sox10*, then we would expect *Sox10* to be less accessible or no longer be accessible in *Nr6a1* null embryos. Indeed, *Nr6a1^-/-^* embryo NCC exhibited decreased chromatin accessibility at *Sox10* compared to controls, consistent with NR6A1 being required for access and transcription of *Sox10* (Figure 5B). The DR0 sites overlapped with the *Sox10* peaks as well, supporting the idea that NR6A1 binds to and regulates the transcription of *Sox10* (Figure 5B).

**Figure 5.**
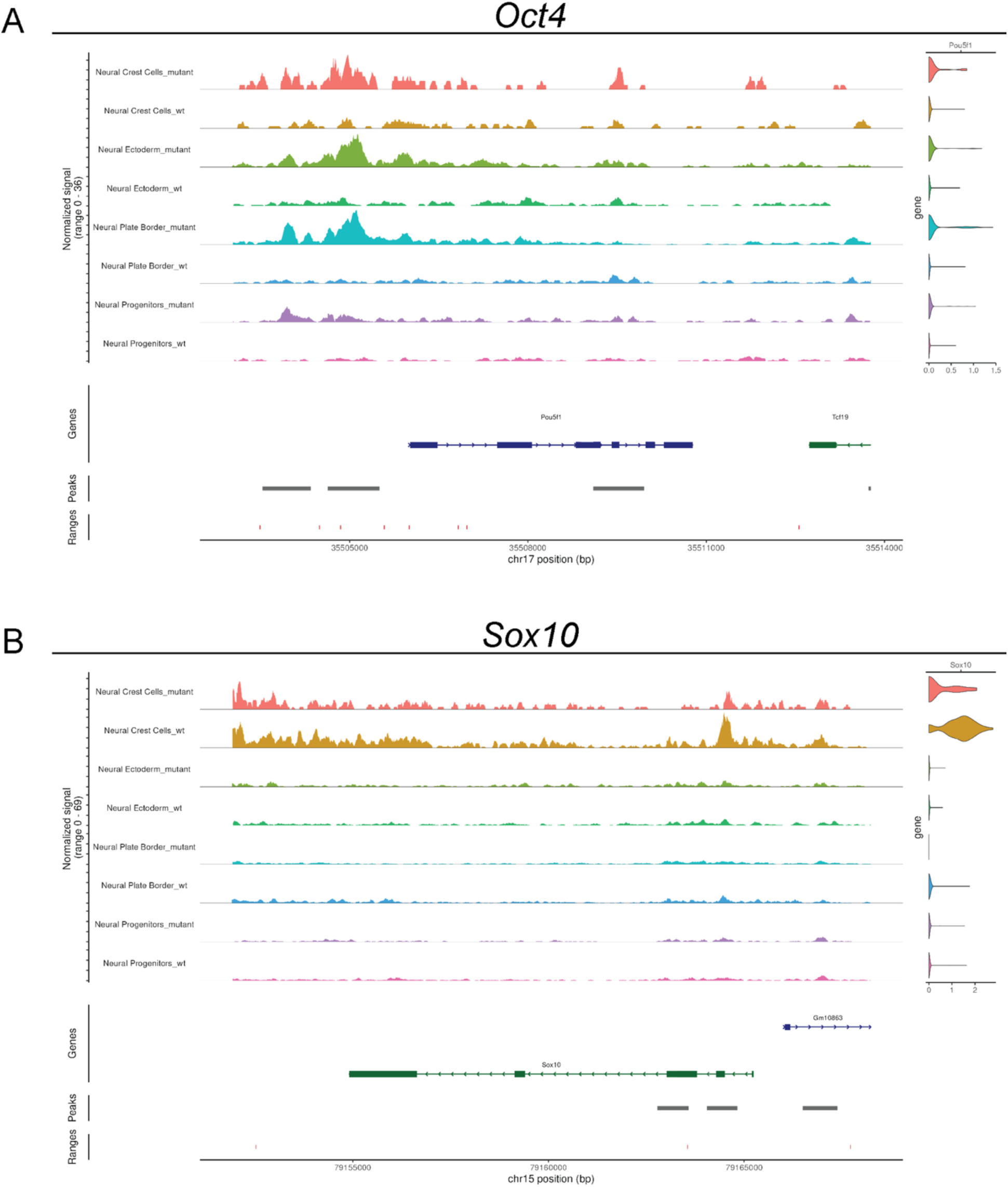
Multiomic scRNA-seq and snATAC-seq demonstrate chromatin accessibility changes around pluripotency-associated and NCC-specifier genes within the NCC and neural plate populations upon loss of NR6A1 function. For both **A** and **B**, the snATAC-seq fragment density across each gene is shown alongside a violin plot displaying gene expression within each cell population. A map of the gene is shown below the fragment density to provide locational reference. Peak coverage is displayed below the gene and the ranges below the peak coverage label *Nr6a1* DR0 binding sites along the genome. A. *Oct4* expression is increased in mutant cell populations compared to wild type as displayed by the violin plot. Consistent with the increased gene expression in the mutant cell populations, the fragment density if higher along *Oct4* indicating the chromatin is more accessible along the gene. Finally, peak coverage overlaps with footprints of DR0 sites which aligns with the requirement of NR6A1 to downregulate *Oct4* expression. B. *Sox10* expression is decreased in the mutant NCC population compared to wild type cells as displayed by the violin plot and matches our previous data showing *Sox10* expression is downregulated in the null. Note that this is normalized to account for fewer NCC in the mutant. When comparing the density of fragments observed along *Sox10* in the snATAC-seq data, there is a reduction in density in the null compared to control NCC. Further, peak coverage overlaps with DR0 bindings sites. Collectively, the data shows there is a reduction in chromatin accessibility along *Sox10* in the null NCC compared to wild type which translates to decreased expression. It is possible this is due to a role of NR6A1 as a regulator of *Sox10* expression according to the overlap in peak coverage and DR0 binding sites. Due to the loss of NR6A1 function, NR6A1 cannot bind to the DR0 sites along *Sox10* in the null thus the chromatin remains inaccessible for transcription.

ATAC-seq demonstrated that NR6A1 can modulate the accessibility of pluripotency-associated and NCC-specifier genes like *Oct4* and *Sox10* respectively (Figure 5). To validate NR6A1’s ability to directly bind to the DR0 binding sites we performed electrophoretic mobility shift assays (EMSA) and competitive inhibition assays. EMSA revealed the presence of shifted *Zeb2* and *Sox10* bands following the addition of NR6A1, indicating NR6A1 can directly bind to the DR0 sites in their promoter regions (Figure 6A). For the competitive inhibition assay, wildtype and mutated DR0 site DNA were introduced in increasing amounts (Figure 6B). We observed diminished binding in the presence of increasing wildtype DNA compared to mutated DR0 site DNA, supporting the ability of NR6A1 to selectively bind to the DR0 sites of *Zeb2* and *Sox10* (Figure 6B). To confirm Nr6a1 binding occurs *in vivo* and to evaluate all of our NCC and EMT genes of interest, we performed a targeted ChIP assay using mESC that were differentiated into NCC. NR6A1 bound the DR0 sites in *Sox9*, *Sox10*, *Snai1* and *Zeb2* (Figure 6C). We also confirmed NR6A1 that can bind *Nanog* DR0 sites, which had previously been shown (Gu, LeMenuet et al. 2005), further supporting our results (Figure 6C).

**Figure 6.**
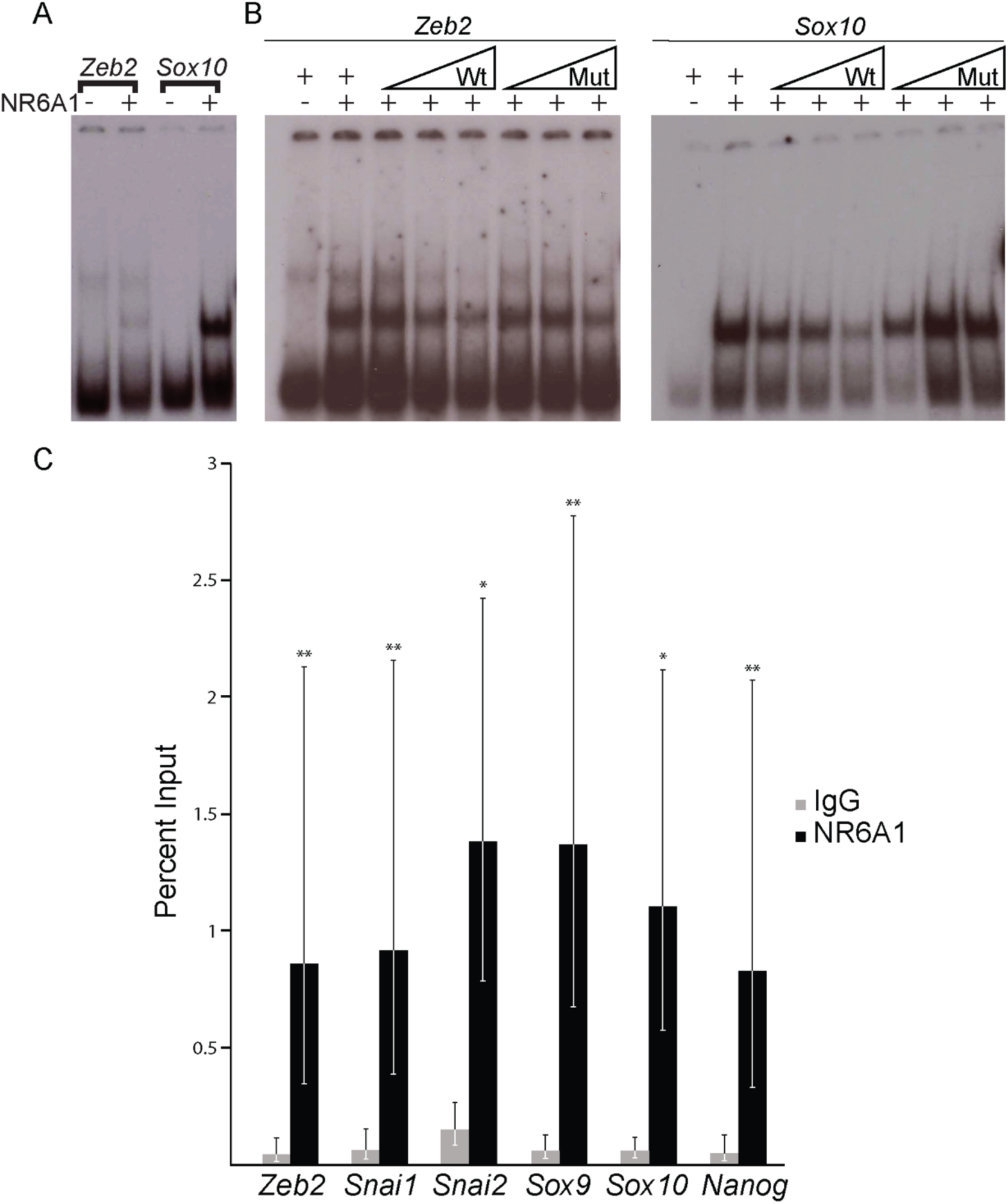
NR6A1 directly binds to putative DR0 binding sites in NCC-specifier and EMT genes. **A.** The ability for NR6A1 to bind to predicted sites of regulation on *Zeb2* and *Sox10* was analyzed by EMSA. When NR6A1 was present, a super-shifted band can be seen, representing positive interactions between NR6A1 and the predicted DR0 sequence. **B.** Unlabeled wildtype sequences or sequences with mutated NR6A1 binding sites for *Zeb2* and *Sox10* were added in a dose-dependent manner resulting in reduced binding to labelled sequences in wildtype and minimal effect with mutant sequence. **C.** A targeted ChIP assay was conducted in mESCs differentiated to NCC using antibodies against endogenous NR6A1 and binding was assessed by qPCR of predicted binding sites. NR6A1 binds to NCC-specifier and EMT factors *in vivo* (*p<0.0, **p<0.005).

In conclusion, NR6A1 can modulate chromatin accessibility and bind to the DR0 sites found in the promoter regions of NCC-specifier and EMT genes, and pluripotency associated factors. The downregulation of NCC-specifier and EMT genes in *Nr6a1* null embryos indicates NR6A1 may function as a transcriptional regulator of NCC-specifier and EMT genes. At the same time, Nr6a1 transcriptionally represses pluripotency associated factors. Collectively these results indicate that Nr6a1 governs the transition of neural stem cells to NCC, by repressing stem cell maintenance and proliferation in the neural ectoderm, while concomitantly regulating the expression of NCC-specifier and EMT genes within the neural plate border and newly emigrating NCC.

### Temporal and tissue-specific excision of Nr6a1 reveals NCC specification begins at mid-late gastrulation

*Nr6a1* null embryos are embryonic lethal by E10.5 due to the failure of chorioallantoic fusion (Chung, Katz et al. 2001). This early lethality prevents the assessment of a role for *Nr6a1* in later NCC differentiation at all axial levels. However, given the important role of Nr6a1 in NCC formation, we posited that Nr6a1 should therefore be essential for craniofacial, cardiac, peripheral nervous system, and gastrointestinal development, as each of these tissues and organs depend heavily on NCC for their proper development. To bypass the early embryonic lethality of Nr6a1 null embryos, we conditionally deleted NR6A1 specifically in the NCC, using *Wnt1-Cre*. Surprisingly, no phenotypic abnormalities were found in *Nr6a1^fx/fx^;Wnt1-Cre* embryos compared to controls (Supplemental Figure 4). E18.5 *Nr6a1^fx/fx^;Wnt1-Cre* embryos appeared morphologically normal, and the NCC derived craniofacial skeleton (evaluated by alizarin red and alcian blue staining), peripheral nervous system and enteric nervous system (evaluated by TUJ1 immunostaining) were indistinguishable from controls. Furthermore, the NCC-specific knockouts were produced in Mendelian ratios and survived until adulthood with full reproductive capabilities. This indicated that NCC development was unaffected (Supplemental Figure 4), which is in stark contrast to all of our other data clearly showing that *Nr6a1* is required for NCC specification, formation and survival.

Recently, however, it was proposed that *Wnt1-Cre* excision occurs too late during neurulation to properly asses gene function during NCC specification and formation in mouse embryos (Barriga 2015). *Wnt1-Cre* facilitates Cre excision of floxed alleles at approximately E8.5. This is important because it suggests that Nr6a1 may not be required for NCC development after E8.5. Furthermore, it also implies that Nr6a1 may instead be needed earlier in development to elicit its role in NCC specification and formation

To test this idea, and establish when Nr6a1 is specifically required, we globally deleted NR6A1 in a temporally-specific manner using the tamoxifen inducible Cre line, *ERT2-Cre* (Indra, Warot et al. 1999). Consistent with the *Wnt1-Cre* knockout, no morphological differences were observed when *Nr6a1* was temporally excised with *ERT2-Cre* at E8.5. We then excised *Nr6a1* via tamoxifen administration at progressively earlier half-day time points. Global excision at E7.75 or E7.25 resulted in no observable phenotypic differences in NCC gene expression in conditional *Nr6a1* mutants compared to controls (Figure 7). However, we discovered that global excision of *Nr6a1* by E6.75 resulted in a phenotype matching the original *Nr6a1* null embryos as evidenced by the downregulated expression of NCC-specifier and EMT genes (Figure 7). Thus, *Nr6a1* is required to induce NCC specification during mid-late gastrulation in mouse embryogenesis, which is considerably earlier than previously appreciated or recognized.

**Figure 7.**
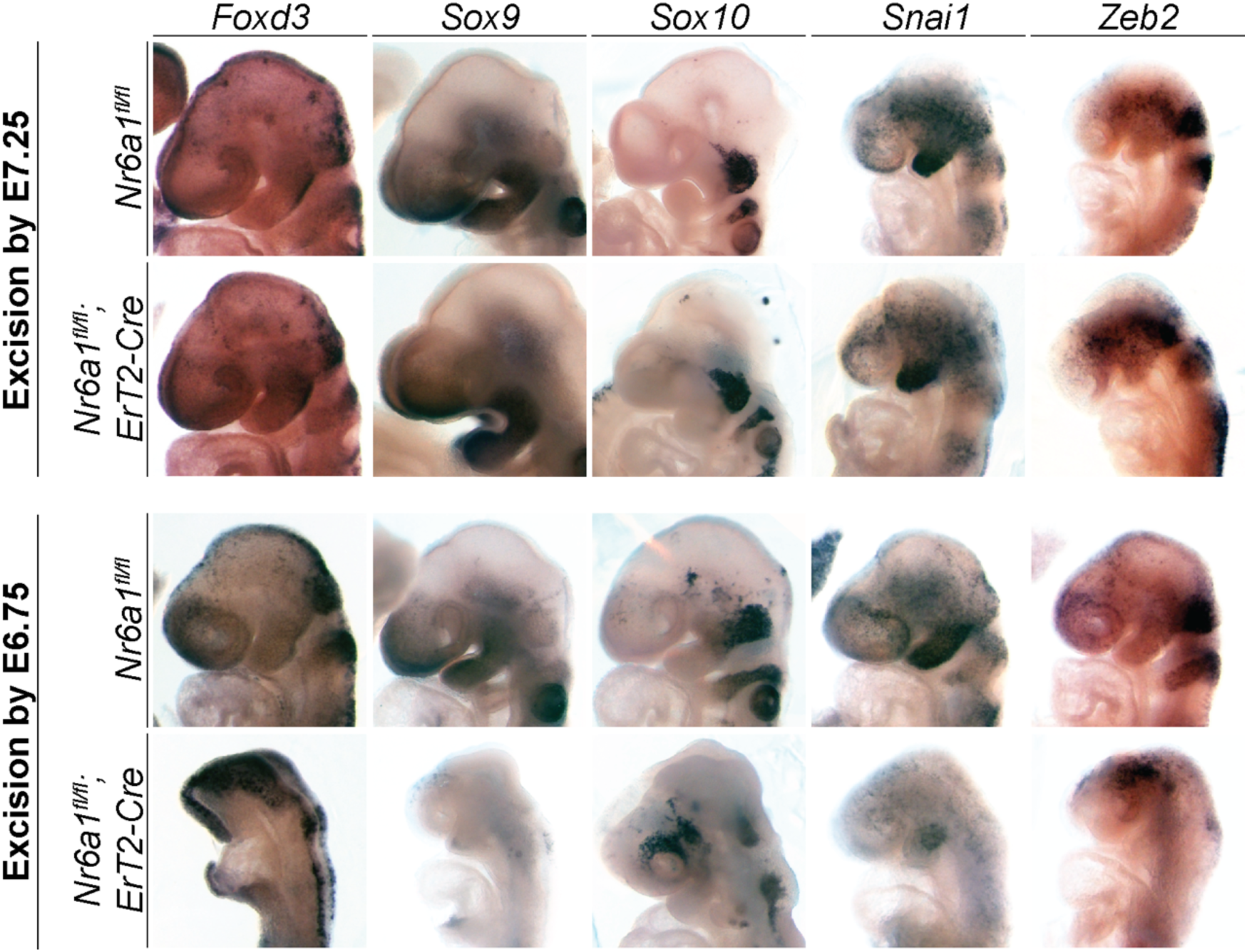
*Nr6a1* activity is required during mid-to-late gastrulation to induce NCC specification. *In situ* hybridization of NCC markers on control (*Nr6a1^fl/fl^*) and *Nr6a1^fl/fl^;ErT2-Cre* embryos treated with tamoxifen for excision of *Nr6a1* at E7.25 and E6.75. Embryos with *Nr6a1* excision by E7.25 and controls show normal expression patterns of NCC and EMT markers. Once *Nr6a1* is excised by E6.75, NCC and EMT marker expression is lost. Auto-color was applied to enhance the contrast above background.

### The downregulation of pluripotency and stem cell maintenance factors is required for NCC formation

Interestingly, it was recently proposed that NCC retain or re-activate pluripotency-associated genes to facilitate their characteristic capacity to generate a wide variety of cell and tissue derivatives (Buitrago-Delgado, Nordin et al. 2015, Lignell, Kerosuo et al. 2017, Zalc, Sinha et al. 2021, Hovland, Bhattacharya et al. 2022, Patel and Parchem 2022, Pajanoja, Hsin et al. 2023). However, our analyses of *Nr6a1^-/-^* mutant embryos indicate that the expression of pluripotency-associated factors including *Oct4*, *Nanog* and *Sox2*, persists in *Nr6a1^-/-^* mutant embryos, coinciding with deficient NCC generation. This suggests that retention of pluripotency-associated factors may actually restrict the generation of NCC from neural stem cells. Consistent with this idea, our analyses of wildtype Oct4-EGFP mouse embryos revealed that Oct4-EGFP expression in E7.5 mouse embryos is downregulated by E8.5, coinciding with the formation of NCC (Figure 4). In further support of these findings, our multiomic analyses revealed a similar absence of *Oct4* and *Nanog* expression in E8.75 wildtype mouse embryos (Supplemental Figure 2A).

To better substantiate whether *Oct4* and other pluripotency and stem cell maintenance factors are downregulated in concert with NCC formation, we analyzed our scRNA-seq data of cranial tissues from pre-migratory and migratory NCC stage mouse embryos (Falcon, Watt et al. 2022, Moore, Zhao et al. 2024, Zhao, Moore et al. 2024). *Sox2, Sox1* and *Nr6a1* are highly expressed in the neural ectoderm, in contrast to *Oct4* and *Nanog* (Figure 4C, Supplemental Figure 5). Furthermore, the expression of each of these genes is downregulated in the progenitor NCC population (Figure 4C, Supplemental Figure 5). In contrast, NCC-specification (*Foxd3*, *Sox9*, *Sox10*) and EMT (*Snai1*, *Zeb2*) genes are highly enriched in progenitor NCC as compared to the neural ectoderm, distinguishing these two populations (Figure 4C, Supplemental Figure 5). Other groups have reported a similar absence of pluripotency-associated factor expression during NCC development aligning with our data (Briggs, Weinreb et al. 2018). Therefore, to expand and further substantiate our assessment of the dynamics of *Oct4*, *Nanog* and *Sox2* expression during development, we also analyzed a publicly available scRNA-seq dataset that covered gastrulation to neurulation stage in mouse embryos (Pijuan-Sala, Griffiths et al. 2019). Complementary to our data, *Oct4*, *Nanog* and *Sox2* were absent from the progenitor NCC cluster, and a definitive boundary was evident in the pseudotime trajectory separating the primitive neuroectoderm from progenitor NCC (Supplemental Figure 6). *Nr6a1* expression in this dataset aligned with our in situ data confirming that *Nr6a1* is expressed in the epiblast during gastrulation, before becoming enriched in the neural ectoderm during neurulation (Supplemental Figure 6).

To further evaluate pluripotency and stem cell maintenance factor expression during NCC formation, we measured by RT-qPCR, the dynamics of *Oct4*, *Nanog*, *Sox2*, *Sox9* and *Sox10* expression during mESC and hiPSC derivation of NCC (Supplemental Figure 7A and C). Each approach revealed the absence of pluripotency-associated factor expression in the forming NCC populations, and this was strikingly exemplified by their immediate downregulation in hiPSC upon provision of NCC differentiation media (Supplemental Figure 7A and C). Interestingly, *Nr6a1* was also specifically elevated (days 1 and 2) in hiPSC during the switch between pluripotent stem cell maintenance, and NCC specification (Supplemental Figure 7C). Immunostaining further substantiated the downregulation in expression of pluripotency-associated factors and upregulation of NCC-specifier genes (Supplemental Figure 7B and D).

To conclusively show that downregulation of pluripotency and stem cell maintenance regulators is required for NCC formation, we overexpressed *Oct4* in mouse embryos from E6.5 to E8.5 (during NCC specification and formation) using a doxycycline inducible system (Hochedlinger, Yamada et al. 2005) (Figure 4D). The temporal initiation of *Oct4* overexpression was designed to correlate with the temporal requirement for Nr6a1. Embryos overexpressing *Oct4* presented with neural tube closure defects and craniofacial abnormalities, mimicking the *Nr6a1* null phenotype. Moreover, the expression of NCC-specifier (*Foxd3*, *Sox9*, *Sox10*) and EMT (*Zeb2*, *Snai1*) genes in these *Oct4* overexpressing embryos was downregulated, similar to that in *Nr6a1* null mutant embryos (Figure 4D). Downregulation of pluripotency and stem cell maintenance factor expression is therefore required for proper NCC formation. Altogether, our data suggests that NR6A1 is a key regulator of the transition of neural stem cells to NCC by repressing pluripotency-associated genes and regulating the expression of NCC-specification and EMT genes during mid-late gastrulation.

## Discussion

Despite decades of research, we have a poor understanding of the mechanisms, signals and gene regulatory networks that regulate mammalian NCC formation. In fact, to date, very few factors essential for mammalian NCC formation have been identified. Here, we demonstrate that NR6A1 is essential for NCC formation as a novel regulator of the transition of neural stem cells to NCC. *Nr6a1* is dynamically expressed in the neural ectoderm and newly emigrating NCC, and NR6A1 loss-of-function results in a deficiency of anterior cranial NCC, and an absence of migrating NCC caudal to the first pharyngeal arch. This cellular phenotype occurs in association with the downregulated expression of NCC specifier (*Foxd3*, *Sox9*, *Sox10*) and EMT (*Snai1*, *Zeb2*) genes, and the persistent expression of pluripotency and stem cell maintenance factors (*Sox2*, *Oct4*, *Nanog*). Analysis of NR6A1 binding to DR0 sites in the promoter regions of NCC-specifier and EMT genes and pluripotency associated genes, correlated with changes in chromatin accessibility, suggesting that NR6A1 directly regulates the activity of these factors. Thus, we propose that NR6A1 regulates murine NCC formation by repressing pluripotency and stem cell maintenance gene expression, while concomitantly inducing NCC specification and EMT genes, which collectively facilitates the transition of neural stem cells to neural crest cells.

Our evaluation of *Nr6a1* in NCC development through tissue-specific and temporally restricted excision allowed us to determine when Nr6a1 was precisely required to elicit its functions. Although *Nr6a1* null mutant embryos exhibit clear defects in NCC development, conditional excision of *Nr6a1,* specifically in pre-migratory NCC using the gold standard *Wnt1-Cre* transgenic line, did not result in an observable phenotype. This was surprising, but it has been posited that *Wnt1-Cre* excision occurs too late to asses gene function during NCC specification and formation in mouse embryos (Barriga 2015). Our work provides some of the first evidence in accord with this idea, since global temporal deletion revealed that Nr6a1 is precisely required by E6.75 to replicate the original Nr6a1 null mutant phenotype. Interestingly, the onset of NCC specification was classically considered to occur during neurulation, after establishment of the three germ layers and subsequent subdivision of the ectoderm into neural ectoderm and non-neural ectoderm (Selleck and Bronner-Fraser 1995). However, recent studies have suggested that NCC induction and specification begin earlier, during, or even preceding gastrulation (Basch and Bronner-Fraser 2006, Steventon, Araya et al. 2009, Buitrago-Delgado, Nordin et al. 2015, Betters, Charney et al. 2018, Prasad, Charney et al. 2020). Similar to these studies in avian and frog embryos (Basch and Bronner-Fraser 2006, Steventon, Araya et al. 2009), our results imply that murine NCC specification also begins during mid-gastrulation. In support of this model and the temporal role of Nr6a1, it’s important to note that Nr6a1 has also been proposed to regulate the transition from primitive to definitive neural stem cells in the neural ectoderm during gastrulation (Akamatsu, DeVeale et al. 2009).

The ability of NR6A1 to act as a transcriptional repressor of pluripotency-associated factors and activator of NCC-specifiers, combined with the deficiency of NCC in *Nr6a1* mutants, supports a model in which pluripotency and stem cell maintenance regulators must be downregulated for NCC formation. However, some recently published studies have argued that NCC either retain or re-acquire the expression of pluripotency-associated factors such as *Oct4* and that this underpins their formation, multipotency and other properties *(Buitrago-Delgado, Nordin et al. 2015, Lignell, Kerosuo et al. 2017, Zalc, Sinha et al. 2021, Hovland, Bhattacharya et al. 2022, Patel and Parchem 2022, Pajanoja, Hsin et al. 2023)*. For instance, when *Oct4* was knocked out in mouse embryos at E7.5 this resulted in a smaller frontonasal mass which was interpreted to mean that *Oct4* is required for ectomesenchyme differentiation of NCC (Zalc, Sinha et al. 2021). However, the expression of *AP2* and *Sox10* was not affected and changes in proliferation were not evaluated despite the global loss of Oct4. It’s also important to note that conditional knockout of Oct4 in the epiblast during gastrulation affected EMT which could impact the generation of NCC (Mulas et al Nichols 2018). Our analyses of *Nr6a1* null embryos challenge the reacquisition model and role for Oct4 in NCC specification and formation. We evaluated the expression of *Oct4* during normal development using the *Oct4-EGFP* transgenic mouse line, which revealed that although *Oct4* is strongly expressed in the epiblast at E7.5, it is downregulated by E8.5 coinciding with the specification and formation of NCC. Furthermore, *Oct4* expression persists ectopically in the neural ectoderm of *Nr6a1*^-/-^*;Oct4-EGFP* embryos in concert with deficient NCC formation. Subsequent evaluation of pluripotency-associated gene and stem cell maintenance factor expression during NCC development through analyses of our own and publicly available scRNA-seq datasets, each revealed a similar downregulation of *Oct4* and *Nanog* expression in concert with NCC formation. Notably, by utilizing both *Wnt1-Cre* and *Mef2c-F10N-LacZ* embryos in our scRNA-seq, we were able to finely cluster the NCC populations based on the expression of one or both markers (Zhao, Moore et al. 2024). *Wnt1-Cre* demarcates dorsal neural ectoderm, pre-migratory and migratory NCC, whereas *Mef2c-F10N-LacZ* demarcates only migratory NCC. (Chai, Jiang et al. 2000, Aoto, Sandell et al. 2015). Our scRNA-seq results matched those obtained in *Xenopus* where even when clustering with a bias towards re-activation of pluripotency, presumptive NCC do not express these factors (Briggs, Weinreb et al. 2018). Collectively, this data argues against the need for the retention of pluripotency-associated factor expression for proper NCC development.

To conclusively test the idea that downregulation of pluripotency-associated factor expression is required for NCC development, we overexpressed *Oct4* in mouse embryos from E6.5-E8.5 coinciding with the specification and formation of NCC. This perturbed NCC formation, mimicking the *Nr6a1* null mutant phenotype. Thus, persistent *Oct4* expression inhibits NCC formation. However, although Oct4, Nanog and Sox2 comprise a core regulatory complex that maintains the pluripotency in various stem cells, it’s important to note that that simply overexpressing *Sox2* in the neuroepithelium of avian embryos represses NCC formation (Wakamatsu, Endo et al. 2004, Remboutsika, Elkouris et al. 2011). Furthermore, neurospheres selected for high levels of Sox2 expression fail to generate NCC upon transplantation into the neuroepithelium of chicken embryos (Remboutsika, Elkouris et al. 2011). Additionally, expressing *Oct4* and *Sox2* in late migratory NCC disrupts their differentiation (Hovland, Bhattacharya et al. 2022). Collectively this corroborates the requirement to downregulate pluripotency-associated factors for proper NCC development, and our data indicates that Nr6a1 is a key regulator of this process.

The duality of NR6A1 to both downregulate pluripotency-associated factors and activate NCC-specification and EMT genes suggest it functions as a bi-modal switch. However, future studies will be necessary to determine how exactly NR6A1 activates NCC-specification gene expression and whether it is truly direct. NR6A1 lacks the canonical activation function-2 (AF-2) domain which bestows activation activity to nuclear receptors in the superfamily (Greschik and Schule 1998, Hummelke and Cooney 2001). However, mutations in the H12 region of *Nr6a1*, where the AF2 domain is typically located in other nuclear receptors, demonstrated the ligand binding domain remains capable of undergoing conformational changes consistent with switching between activator and repressor functions (Greschik, Wurtz et al. 1999, Hummelke and Cooney 2001). Identification of NR6A1’s ligand with subsequent assays to assess direct transcriptional upregulation will help elucidate whether NR6A1 is indeed a bi-modal switch capable of repression and activation like its most homologous nuclear receptor superfamily member, COUP-TF.

Regardless of whether NR6A1 directly regulates NCC-specifier and EMT genes, we have rigorously demonstrated that NR6A1 is required for proper NCC specification and formation. Interestingly, although the expression of NCC-specification and EMT genes are largely absent in *Nr6a1* null embryos, lineage tracing revealed that some *Wnt1-Cre*-positive cells delaminate and migrate into the facial prominences and first pharyngeal arch. These putative NCC cells, which are of undefined character, ultimately undergo cell death as evidenced by TUNEL staining, demonstrating that NR6A1 is required for NCC survival. How these cells remain capable of delamination without the expression of EMT master regulators *Snai1* and *Zeb2* remains a mystery. However, it should be noted that individual knockouts of *Snai1* and *Zeb2* and even compound *Snai1/2* double knockouts in mice do not prevent NCC specification and delamination (Van de Putte, Maruhashi et al. 2003, Murray and Gridley 2006, Van de Putte, Francis et al. 2007). In these scenarios, *Snai1* and *Zeb2* have been proposed to compensate for each other to justify the presence of delaminated NCC. However, in *Nr6a1* null embryos, the expression of both of these factors is downregulated, and yet some putative NCC still delaminate from the neural ectoderm and migrate into the facial prominences and first pharyngeal arch. The expression of *Twist1* in these cells indicates they are likely mesenchymal which could explain their migratory ability. However, *Twist1* is not expressed by NCC in mouse embryos until after their delamination from the neural ectoderm, and consistent with this behavior, Twist1 loss-of-function does not affect NCC specification and delamination (Gitelman 1997, Tavares, Izpisuja-Belmonte et al. 2001, Bildsoe, Loebel et al. 2009). Thus, *Twist1* expression in *Nr6a1* null mutants does not explain the ability of *Wnt1-Cre;R26R-eYFP-*positive cells to delaminate from the neural ectoderm. Interestingly, we recently discovered that a subpopulation of NCC can delaminate via a novel cell extrusion mechanism in parallel with canonical EMT (Moore, Zhao et al. 2024). This could explain the presence of delaminated migratory cells in *Nr6a1* mutant embryos, however future experiments are needed to determine if this is indeed the case in addition to identifying specific markers of NCC extrusion.

In summary, our work has demonstrated that pluripotency and stem cell maintenance factors need to be downregulated in association with the activation of NCC-specification and EMT genes, to facilitate the differentiation of neural stem cells to NCC. This transition in signaling is facilitated by NR6A1, which is required during mid-late gastrulation. Altogether, our findings have expanded our knowledge of the regulatory signals and mechanisms driving NCC formation as well as the timing of NCC specification in mammals, deepening our overall understanding of NCC development.

## Methods

### Animal husbandry and embryo collection

All mice were housed in the Laboratory Animal Services Facility at the Stowers Institute for Medical Research under a 14 hour light/10 hour dark, light cycle, and experiments were performed according to IACUC animal welfare guidelines and approved protocol (#2022-014). Embryos were obtained by timed mating, with the morning of the vaginal plug defined as embryonic day (E)0.5 and genotyping of yolk sac tissue was performed as previously described (Chung, Katz et al. 2001, Lan, Xu et al. 2003).

For temporal excision of *Nr6a1*, pregnant dams were orally gavaged with 5 mg of Tamoxifen and 1 mg of progesterone dissolved in 100 µL of corn oil. Treatment was performed one day prior to the desired stage of allele excision as it takes at least 24-hours for complete recombination to occur.

To overexpress *Oct4*, CD1 females were mated with *Oct4/rtTA (Col1a1-tetO-Oct4;R26R-M2rtTA)* transgenic males (Jax stock #006911). Transgene expression was induced from E6.5 until the desired stage by replacing drinking water of pregnant females with a 7.5% sucrose solution containing 0.5 mg/mL doxycycline. Embryos from non-treated mice of the same genotype were used as controls.

### In situ hybridization

Following dissection, embryos were fixed in 4% paraformaldehyde overnight at 4°C. *In situ* hybridization was then performed with anti-sense digoxigenin-labeled mRNA riboprobes as previously described (Nagy 2003).

### Immunohistochemistry

For section immunohistochemistry, *Nr6a1* litters were collected at E9.0 and fixed overnight in 1% PFA at 4°C. Embryos were processed through a sucrose gradient, mounted in Tissue Tek OCT (VWR, West Chester, PA) and sectioned at 10 microns. Sections were rinsed 3 times with PBS 0.1% TritionX-100 (PBT) for 5 minutes and blocked in 10% goat serum (Invitrogen, Carlsbad, CA) in PBT for 1 hour at room temperature. Slides were incubated in primary antibody diluted in blocking solution at 4°Covernight. The monoclonal antibodies SOX2 (R & D Systems, Minneapolis, MN) and pHH3 (Upstate/Millipore, Billerica, MA) were both used at 1:500. The antibody to PAX3 was obtained from the Developmental Studies Hybridoma Bank (Iowa City, IA 52242). TUNEL staining (Roche Life Sciences) was performed according to the manufacturer’s instructions. Slides were rinsed 3 times for 10 minutes in PBT at room temperature on a shaker and incubated in the appropriate Alexa secondary antibody (Molecular probes/Invitrogen, Carlsbad, CA) at 1:250 for 2 hours at 4°C. Sections were counterstained with a 1:1000 dilution of 2mg/mL DAPI (Sigma, St. Louis, MO) in PBS for 5 minutes, followed by rinses in PBS before mounting slides with VectaShield (Vector Laboratories, Newark, CA). All images were collected using a Zeiss Axioplan microscope and processed using Photoshop CS2 (Adobe, San Jose, CA).

Immunostaining of whole embryos and cultured cells was performed as previously described (Barlow, Dixon et al. 2012, Leung, Murdoch et al. 2016). The WTC-11 human induced pluripotent stem cell line, was obtained from the Coriell Institute (#GM25256). hiPSCs were plated in an iBidi 8-well plate (Ibidi #80826), and maintained as pluripotent stem cells or differentiated into neural crest cells. Cells were fixed in 4% PFA (Alfa Aesar via VWR Cat. No. AA43368-9M) for 15 minutes at room temperature and then permeabilized in PBS + 0.3% Tx-100 (Millipore Sigma, #TX1568, PBST) for 20 minutes at room temperature. Cells were blocked using PBST + 0.5% BSA (Millipore Sigma, #2910) for 1-2 hours at room temperature, and incubated in primary antibody in blocking buffer overnight at 4°C. The following primary antibodies were used: Oct4 (R&D Systems, #MAB1759 1:50), Nanog (R&D Systems, #1997 1:40), Sox10 (R&D Systems, #2864 1:100), and Tfap2a (Invitrogen, #3B5 1:50). The cells were washed 3 times for 5 minutes with PBST and 0.5% BSA(Millipore Sigma, #2910) before incubating in secondary antibody and blocking solution. The following secondary antibodies were used at a 1:500 dilution: Donkey anti-Rabbit IgG (H+L) Alexa Fluor™ 546 (ThermoFisher #A10040), Donkey anti-Goat IgG (H+L) Alexa Fluor™ 488 (ThermoFisher, #A11055), and Donkey anti-Mouse IgG (H+L) Alexa Fluor™ 546 (ThermoFisher, #10036) at room temperature for 1-2 hours. The cells were then washed 3 times for 5 minutes with PBST and 0.5% BSA, mounted in Vectashield in DAPI (Vector Labratories, #H-1200-10) and imaged. All images were individually acquired with the same settings, and brightness and contrast were adjusted the same throughout.

### Microarray

E9.0 wild type embryos and *Nr6a1^-/-^* embryos were isolated and total RNA was extracted from whole embryos using TRIzol reagent. Concentration and quality of RNA were determined by spectrophotometer and Agilent bioanalyzer analysis (Agilent Technologies, Inc., Palo Alto, CA). For array analysis, labeled mRNA targets were prepared with 150 ng total RNA using MessageAmp III RNA Amplification Kit (Applied Biosystems / Ambion, Austin, TX) according to the manufacturer’s specifications. Array analysis was performed using Affymetrix GeneChip Mouse Genome 430 2.0 Arrays processed with the GeneChip Fluidics Station 450 and scanned with a GeneChip Scanner 3000 7G using standard protocols. Resulting CEL files were analyzed using RMA and limma in the R statistical environment (Chang, Manent et al. 2022). The microarray data is publicly available in the Gene Expression Omnibus using accession number GSE166458.

### Bulk RNA-sequencing

E8.75 *Nr6a1^-/-^* and wild type control littermates of between 9-11 somites were isolated and bisected into anterior (rostral to and including the first branchial arch) and posterior (caudal to and including the second branchial arch) samples. Total RNA was extracted from whole embryos using an RNA micro Kit. Concentration and quality of RNA was determined by bioanalyzer analysis. RNA-seq was run using single-end 50bp sequencing reads with Illumina HiSeq 4000. FASTQ files were aligned with STAR, and EdgeR was used for differential expression analysis between mutant and wild type (Chang, Manent et al. 2022). The RNA-seq data is publicly available in the Gene Expression Omnibus using accession number GSE180427.

### Single-cell RNA sequencing

Single-cell RNA sequencing of E8.5 *Wnt1-Cre;R26R-eYFP* and *Mef2c-F10N-LacZ* embryos, and subsequent bioinformatic analyses were performed as previously described (Falcon, Watt et al. 2022, Moore, Zhao et al. 2024, Zhao, Moore et al. 2024) and the data set is publicly available at the Gene Expression Omnibus using accession number GSE168351. For single cell RNA sequencing gastrulation to neurulation time courses, the data was obtained and images generated from the interactive server: https://marionilab.cruk.cam.ac.uk/MouseGastrulation2018/ described in (Pijuan-Sala, Griffiths et al. 2019).

### *Multiomic* single-cell RNA-sequencing and single-nucleotide ATAC-sequencing

*Nr6a1^-/-^* and wild type control littermates were collected at E8.75. 3-4 embryos were used per genotype across two litters. Nuclei was isolated with the 10X Genomics Nuclei Isolation kit. Following isolation, the quality of nuclei was assessed via bright field imaging and standard QC analysis. A total of 3,200-3,500 nuclei were captured per sample. Libraries were prepared and pooled such that the pooled gene expression and ATAC libraries could then be run on individual flow cells, resulting in 20,000 reads per nuclei for scRNA-seq and 25,000 reads per nuclei for snATAC-seq. Paired end sequencing was performed on an Illumina NextSeq 2000. Raw data was processed with 10X Cell Ranger and analysis was performed in R (v4.4.1) with Seurat (v5) (Hao, Stuart et al. 2024). Data was filtered using standard QC methods: ATAC reads greater than 50,000 and less than 500 were filtered out along with RNA reads greater than 25,000 and less than 1,000. Cells with percent RNA mitochondria reads greater than 10% were filtered out. Finally, cells in which transcription start site enrichment was greater than 1 or nucleosome signal less than 2 were also filtered out. The final dataset analyzed following filtering and SCT normalization (Choudhary and Satija 2022) included 2,987 mutant cells and 2,728 wild type cells. Clustering was performed on the wild type dataset at a resolution of 0.5 using the find nearest neighbor method. Mutant cells were merged with wild type cells and integrated with CCA Integration (Butler, Hoffman et al. 2018) to compare clusters between the datasets. Annotations from the scRNA-seq clustering were used to annotate the snATAC-seq.

### Electrophoretic mobility shift assay (EMSA) and Competition Binding Assay

We produced NR6A1 protein *in vitro* using the TNT translation system (Promega). Oligo DNA 5’ – hydroxyl end labeled with (γ-^32^P) ATP using T4 PNK system was used as a probe. EMSA experiments were performed at 30°Cfor 30 minutes in 10 µL of EMSA buffer (10mM HEPES, pH 7.5; 50mM KCl; 1mM EDTA, 5mM MgCl_2_; 2mM DTT; 10% Glycerol; Poly [dI-dC] 1µg). For competition assays unlabeled wild type or mutated competitor DNA was added to the reaction at 5x, 10x or 50x concentrations relative to labelled probe. The total reactions were directly loaded onto a 5% acrylamide gel. Electrophoresis was carried out at 4°C for 100 minutes at 300V. The WT sequence for the competition assay was AGGTCA and the Mutant sequence was CTTGAC.

### Mouse stem cell to primitive neural crest cell differentiation

KH2 cells were maintained on MEFs in KOSR media. Prior to initiating differentiation, the cells were adapted to feeder free conditions for 1 passage. For differentiation, 12,500 cells/cm^2^ were plated on Matrigel (Sigma-Adlrich) coated plates. Base differentiation media consisted of DMEM/F12 with Glutamax:Neurobasal 1:1 media containing 1X Embryomax, 2-mercaptoethanol, 0.5X N2, 0.5X, B27, and 1X NEAA. bFGF (10ng/mL) (Stemcell technologies) and Heparin solution (0.0002%) (Stemcell technologies) were added to the media for the entirety of differentiation. After the first 4 days of differentiation, Bmp2 (10 ng/mL) was added to the culture until the cells were harvested.

### Human induced pluripotent stem cell to neural crest differentiation

Undifferentiated hiPSCs were maintained and passaged on plates coated with hESC-qualified Matrigel (Corning #354277) in mTeSR1 (Stem Cell Technologies #85850) media supplemented with 1% penicillin/streptomycin (ThermoFisher, #15070063). Cells were passaged approximately every 3–5 days (70–85% confluency) using Accutase (Gibco, #11105–01) to detach cells. Cells were then plated in mTeSR1 + 1% P/S and 10 μM Rock Inhibitor (Y-27632, StemCell Technologies, #72308). For NCC differentiation, hiPSCs were detached with Accutase (Gibco, #11105–01) and resuspended in STEMdiff^TM^ Neural Crest Differentiation media (StemCell Technologies, #08610). Cells were plated at 7,000 cells/cm^2^ in STEMdiff^TM^ Neural Crest Differentiation media supplemented with 1% penicillin/streptomycin (ThermoFisher, #15070063) and 10 μM Rock Inhibitor (Y-27632, StemCell Technologies, #72308) for the first 24 hrs. Media was changed every day. On day 8 of differentiation, cells were harvested for RNA extraction or fixed for immunostaining.

### Chromatin Immunoprecipitation

ChIP experiments were performed as previously described (Smith, Martin-Brown et al. 2010), using mouse KH2 stem cells that were differentiated into to NCC over 5 days These experiments were repeated 5 times for rigor, reproducibility and statistical analysis. Antibodies used were: NR6A1 (Proteintech Ca. No. 12712-1-AP) and IgG control (Proteintech Ca. No. 30000-0-AP). Snai1 promoter sequence:

CAGCGCCCAAAGGTCAGCAGCTCGGGGATG; Snai2 promoter sequence:

AAGCCAAGTCGCCGTAGGTCACCTAGCGGAA; Sox9 Exon 3 sequence:

GTTCCGGCCACCCACGGCCAGGTCACCTACAC; Sox9 3’ UTR sequence:

GCTGTTCCCCGTGGAGGTCAGAGGGGAGAGGTA; Sox10 Exon 2 sequence:

GTGAACTGGGCAAGGTCAAGAAGGAACAGCA; Zeb2 Intron 2 sequence:

CTTGGCTCCAGCAGTGAGGTCAAGCCACAGCC; Zeb2 Intron 2 sequence 2:

AGAGGTCATGTGAACCTCAGAGTCAGGCCCTCG.

### Quantitative PCR

RNA was extracted from WTC-11 hiPSC or differentiated NCC using the Qiagen miRNeasy Micro Kit (Qiagen, #217084) with on-column DNase treatment (Qiagen, #79254). The Superscript III Kit (Invitrogen, 18080051) was used to synthesize cDNA for quantitative RT-PCR (qPCR) using random hexamer primers. qPCR was performed on ABI7000 (Thermo QuantStudio 7) using Perfecta Sybr Green (Quantbio # 95072-250). Primers are listed in Table 2.1. No template controls were run as negative controls. The ΔΔCt method was used to calculate fold change. Student’s t-test and ANOVA were used for statistical analysis and significance was determined based on p < 0.05.

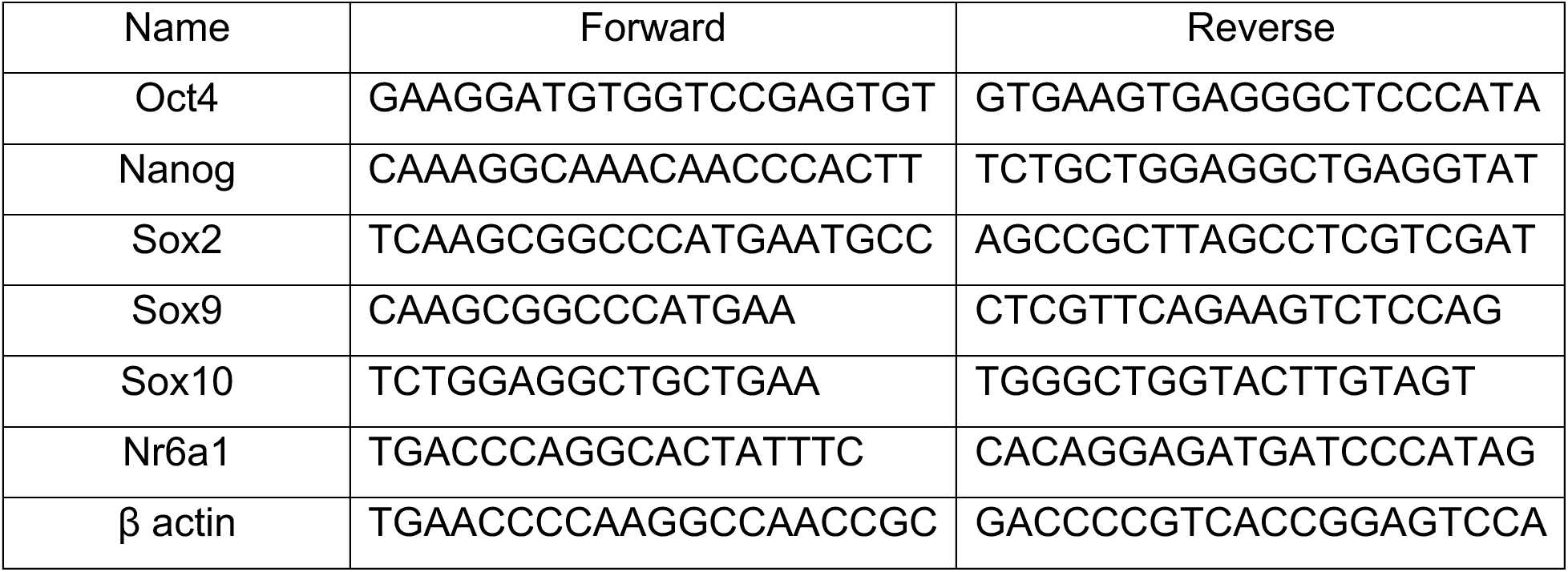

### Imaging and image analysis

Brightfield images of *in situ* hybridization stained embryos were captured as previously described (Zhao, Moore et al. 2024). In brief, images were taken on a Leica MZ16 microscope using a Nikon DS-Ri1 camera. Manual Z-stacks were acquired and then images were merged using Helicon Focus. Fluorescent images were taken on upright Zeiss LSM-700, inverted Zeiss LSM-800 equipped with a GaAsP detector and a Nikon CSU-W1 inverted spinning disk equipped with a sCMOS camera. Image analysis was performed using Fiji.

### Statistical analysis

Results were expressed as a mean with error bars representing standard error of the mean (SEM). Statistical analysis was performed by Student’s t-test, and p-values <0.05 were considered significant.

### Skeletal staining of Wnt1-conditional knockout

E18.5 embryos were anesthetized by immersion in ice cold PBS for at least 60 minutes until no reflex movements were observed following a pinch test. The embryos were then fixed in 95% ethanol for 1 hour and dehydrated in fresh ethanol overnight at 4°C. The skin and viscera were manually removed, and the embryos were incubated in 70% ethanol/5% acetic acid for 30 minutes at room temperature, before staining overnight in alizarin red/alcian blue at room temperature as previously described (Nagy, Gertsenstein et al. 2009, Nagy, Gertsenstein et al. 2009). The embryos were then washed in 70% ethanol/5% acetic acid for 30 minutes and quickly rinsed in water. Soft tissue digestion was performed in 2% KOH overnight for at least 6 hours, followed by further digestion and clearing, initially in 0.25% KOH, and then through an increasing gradient of glycerol in 0.25% KOH before being stored in 50% glycerol.

## Acknowledgements

The authors thank past and present members of the Trainor lab and Manzanares lab for their thoughtful insights, constructive feedback and discussions during the progress of this work. We are extremely grateful to Melissa Childers, Marina Thexton and the Laboratory Animal Services Core for their care and maintenance of our mouse colonies, and we appreciate Kaitlyn Petentler, Allison Scott and the Sequencing Core for their molecular biology support. We especially thank Dr. Austin Cooney for providing the *Nr6a1^+/^-* and *Nr6a1^fx/fx^* mice to establish our own lines. *Nr6a1, Pax3*, *Snai1*, *Sox9*, *Sox10* and *Wnt1* riboprobe plasmids were generously provided by Dr. Austin Cooney, Dr. Takayoshi Inoue, Dr. Angela Nieto, Dr. Trevor Williams, Dr. Martin Gassmann and Dr. Antony Gavalas respectively.

---

**Supplemental Figure 1.**
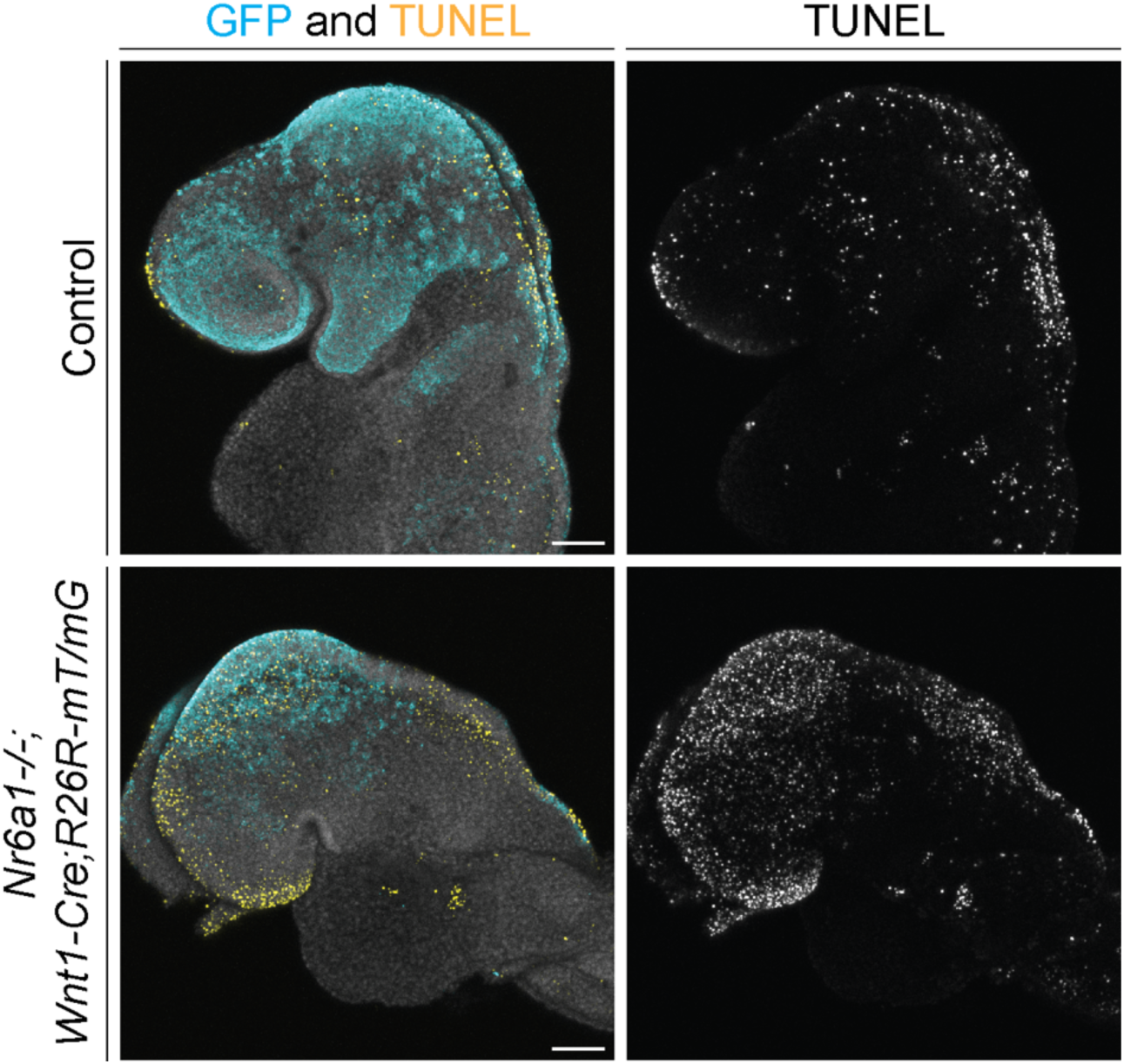
N*r*6a1 is required for NCC survival. TUNEL staining of *Nr6a1^-/-^;Wnt1-Cre;R26R-mT/mG* and *Wnt1-Cre;R26R-mT/mG* control littermates at E8.75 shows high amounts of cell death in the GFP labeled NCC population throughout the head. Therefore, the migratory NCC population in the null embryos do not survive and undergo cell death.

**Supplemental Figure 2.**
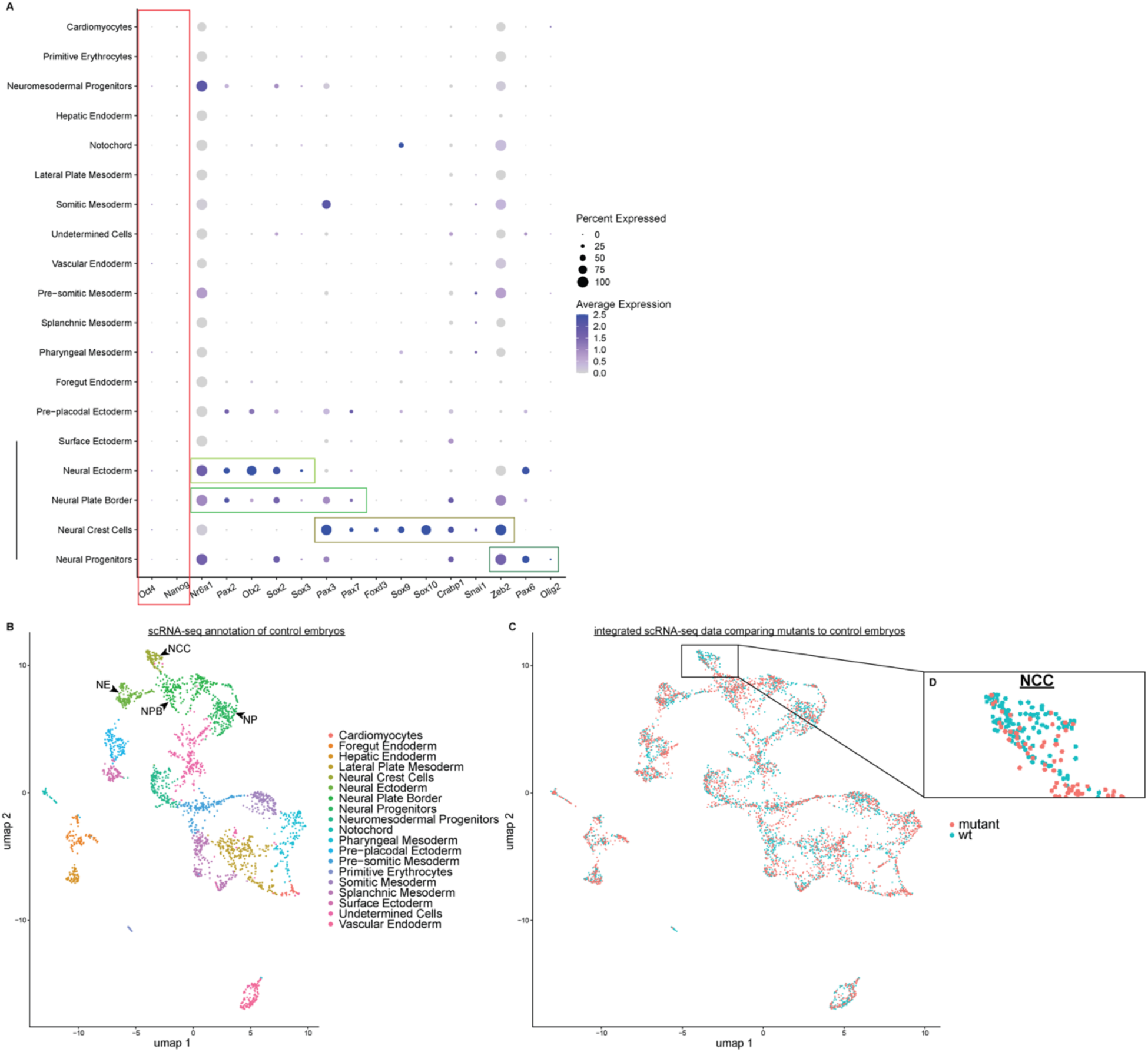
Annotation and integration of scRNA-seq acquired at E8.75-9.0 from *Nr6a1* null and control littermate embryos. **A.** Clustering the control dataset at a resolution of 0.5 resulted in 19 distinct cell populations which were identified based on the expression of canonical marker genes. Notably, *Oct4* and *Nanog* are minimally expressed in the embryo at these stages of development (red box). Neural ectoderm cells were identified by the high expression of *Sox2*, *Pax2* and *Otx2* and low expression of other neural plate genes. Neural plate border cells were identified by the expression of *Sox2*, *Pax2*, *Wnt1*, *Pax3*, and *Pax7* and low expression of NCC genes. NCC were identified by the high expression of NCC genes *Pax3*, *Pax7*, *Foxd3*, *Sox9*, *Sox10*, *Crabp1*, *Snai1* and *Zeb2*. Finally, the neural progenitors were identified based on expression of *Sox2*, *Pax6* and *Olig2*. **B.** UMAP of the fully annotated control scRNA-seq dataset. **C.** UMAP of the *Nr6a1^-/-^* (mutant, red) scRNA-seq dataset integrated with the control scRNA-seq dataset (blue). **D.** Zooming into view the NCC cluster, we can visualize the deficiency of NCC we have consistently observed in the *Nr6a1* null.

**Supplemental Figure 3.**
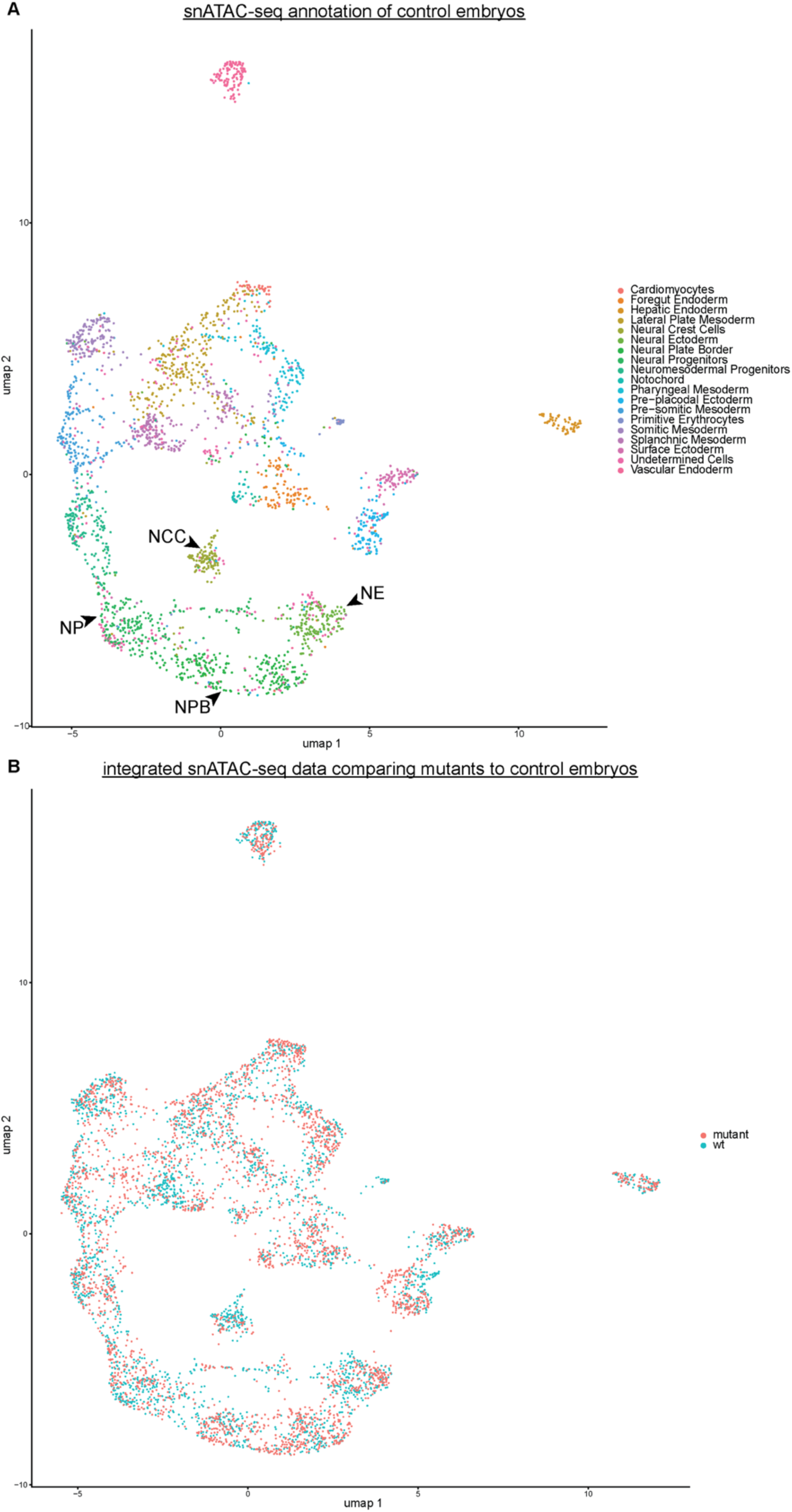
UMAP of the snATAC-seq dataset using the cell population annotations from the scRNA-seq data. **A.** UMAP of the control littermate snATAC-seq dataset displaying the annotated cell populations identified from the scRNA-seq analysis. **B.** UMAP of the *Nr6a1^-/-^* embryo snATAC-seq dataset integrated and overlaid with the control snATAC-seq dataset.

**Supplemental Figure 4.**
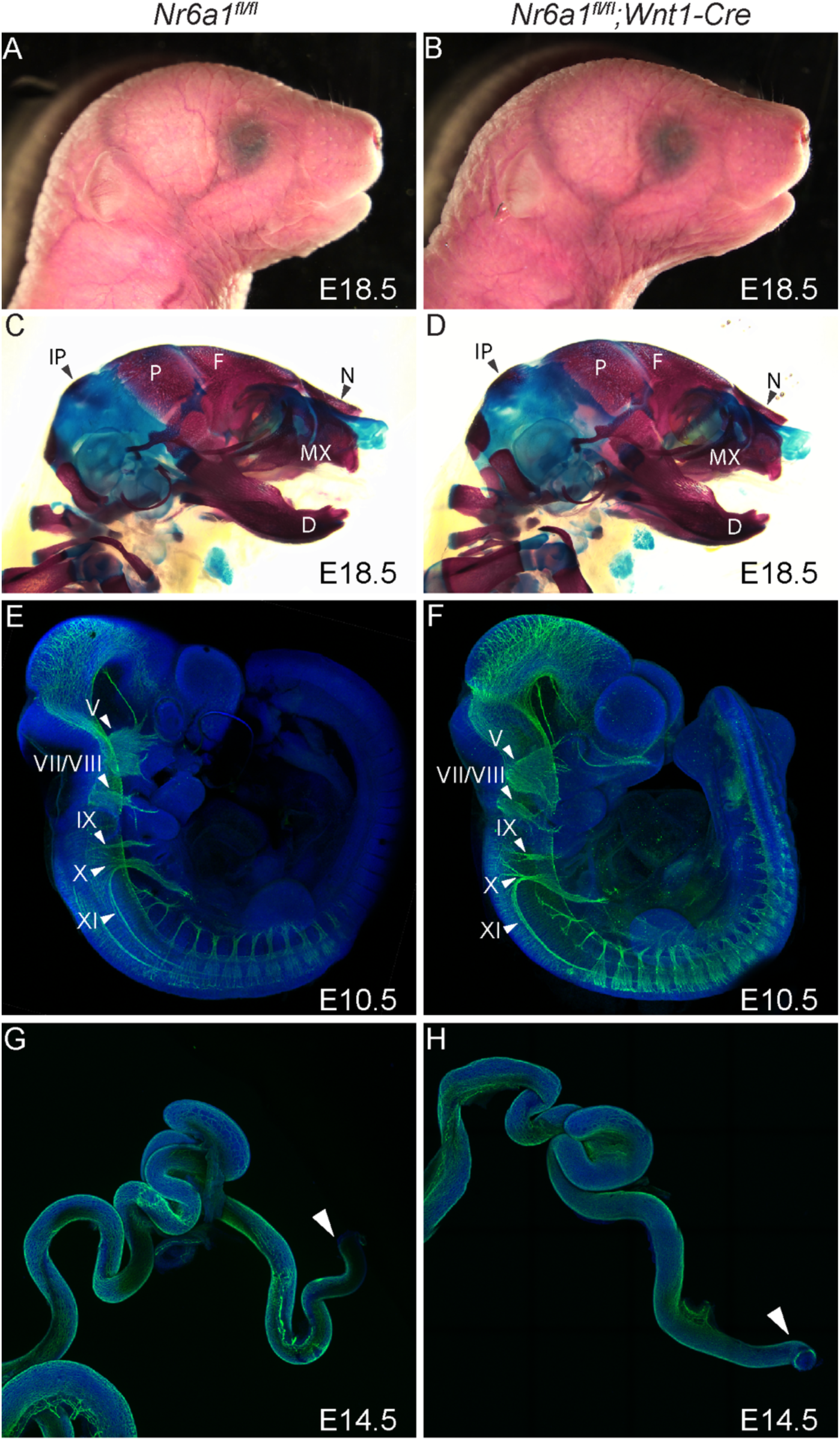
W*n*t1*-Cre* excises *Nr6a1* after it is required for NCC specification and formation. A-B. Brightfield images of *Nr6a1^fl/fl^* and *Nr6a1^fl/fl^;Wnt1-Cre;R26R-eYFP* embryos at E18.5. No differences were observed in the craniofacial morphology. **C-D.** Alcian blue (cartilage) and alizarin red (bone) skeletal staining at E18.5 showed the craniofacial skeleton was normal, supporting our initial assessment of the craniofacial morphology. **E-F.** DAPI (blue) and neurofilament (TUJ1, green) immunostaining of E10.5. embryos indicated no overall changes in the peripheral nervous system. Cranial nerves are labeled with an arrowhead. **G-H.** DAPI (blue) and neurofilament (TUJ1, green) immunostaining of intestines isolated from E14. 5 showed no differences in the pattern of neurofilament formation between the conditional and wildtype embryos. The arrowhead indicates the distal tip of the colon.

**Supplemental Figure 5.**
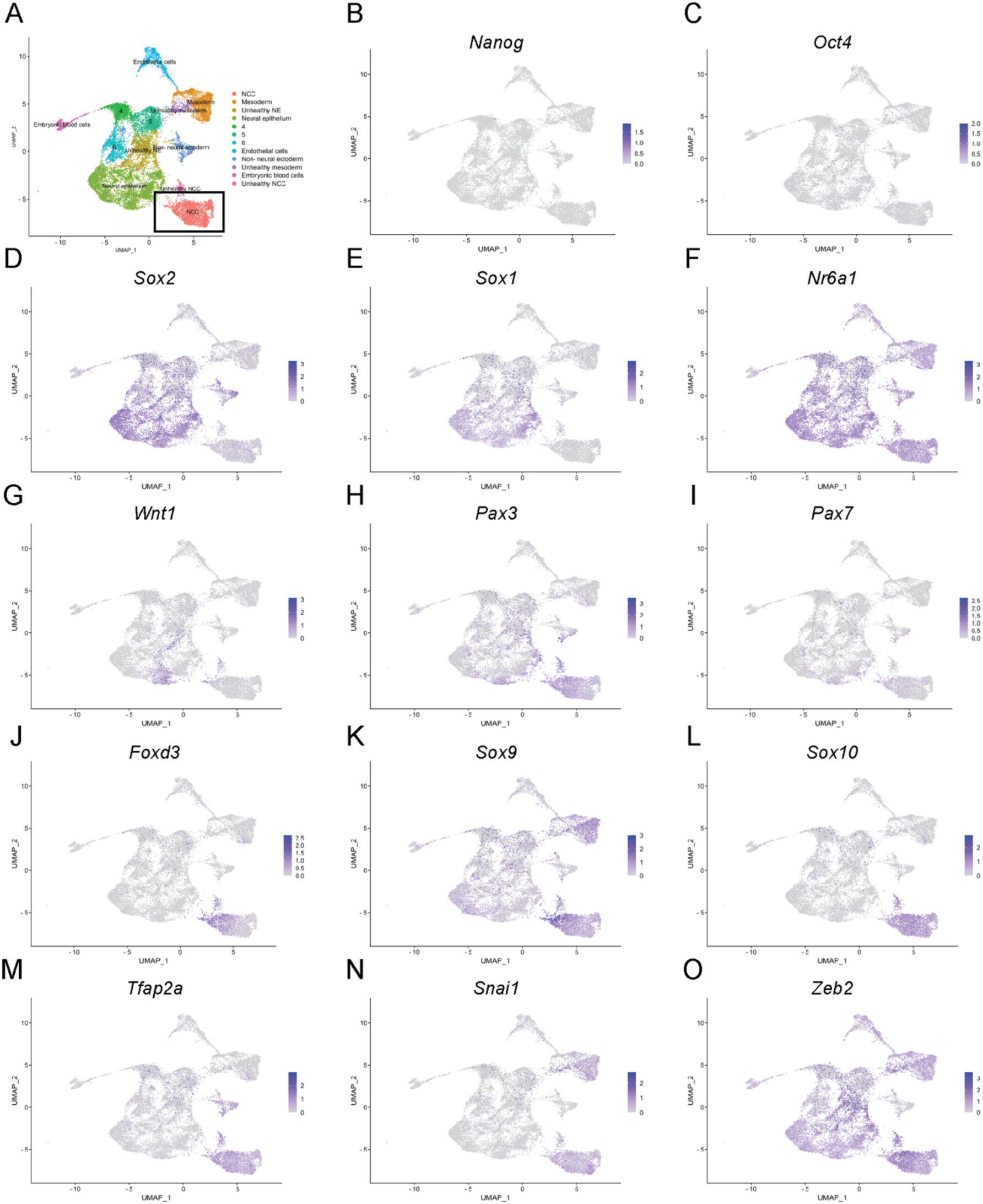
NCC downregulate pluripotency-associated genes prior to undergoing EMT. scRNA-seq of cranial tissue from *Wnt1-Cre;R26R-eYFP* and *Mef2c-F10N-LacZ* embryos covering stages of NCC development. **A.** UMAP plot showing color coded cells annotated by defined cell type. A box outlines the NCC cluster. **B-O.** Expression levels of indicated gene, overlaid on the UMAP of all cell clusters. *Oct4* and *Nanog* are not expressed in the NCC cluster. *Wnt1* expression is restricted to the neural ectoderm while *Pax3* is expressed in the neural ectoderm and early NCC populations. *Foxd3, Sox9, Sox10* and *Tfap2a* are specifically expressed in the NCC cluster.

**Supplemental Figure 6.**
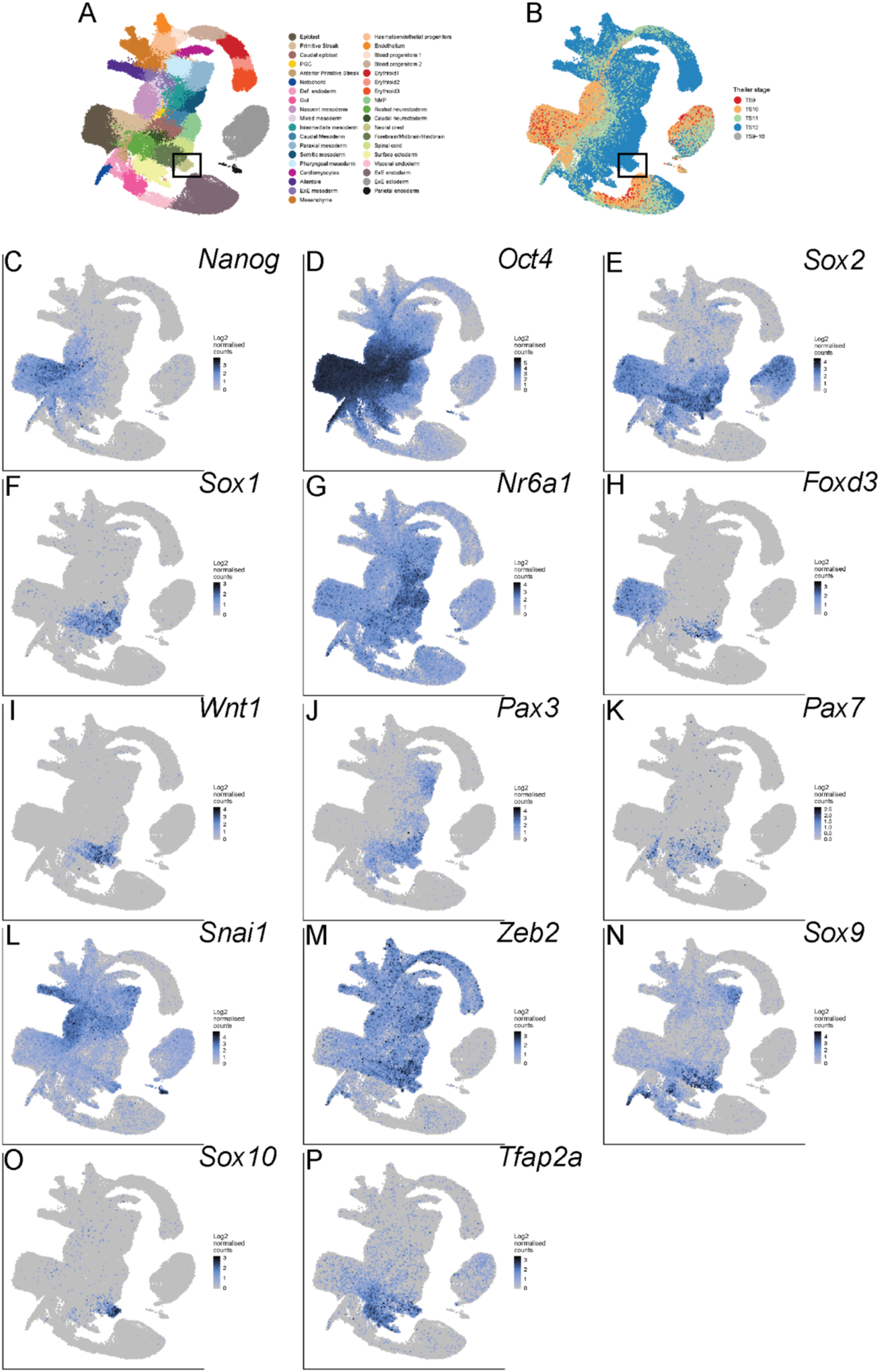
NCC progress through neural ectodermal fate prior to commitment. scRNA-seq data from gastrulation to neurulation stages of development in mouse (Pijuan-Sala, Griffiths et al. 2019). **A.** UMAP plot showing color coded cells annotated by defined cell type. **B.** UMAP plot showing stage profiles color coded based on Theiler stage at time of collection. The box outlines the NCC population in both plots. **C-P.** Expression levels of indicated genes overlaid on the UMAP of all cell types. *Oct4* and *Nanog* are absent from the NCC cluster where expression of NCC-specifiers *Sox9*, *Sox10* and *Foxd3* are enriched.

**Supplemental Figure 7.**
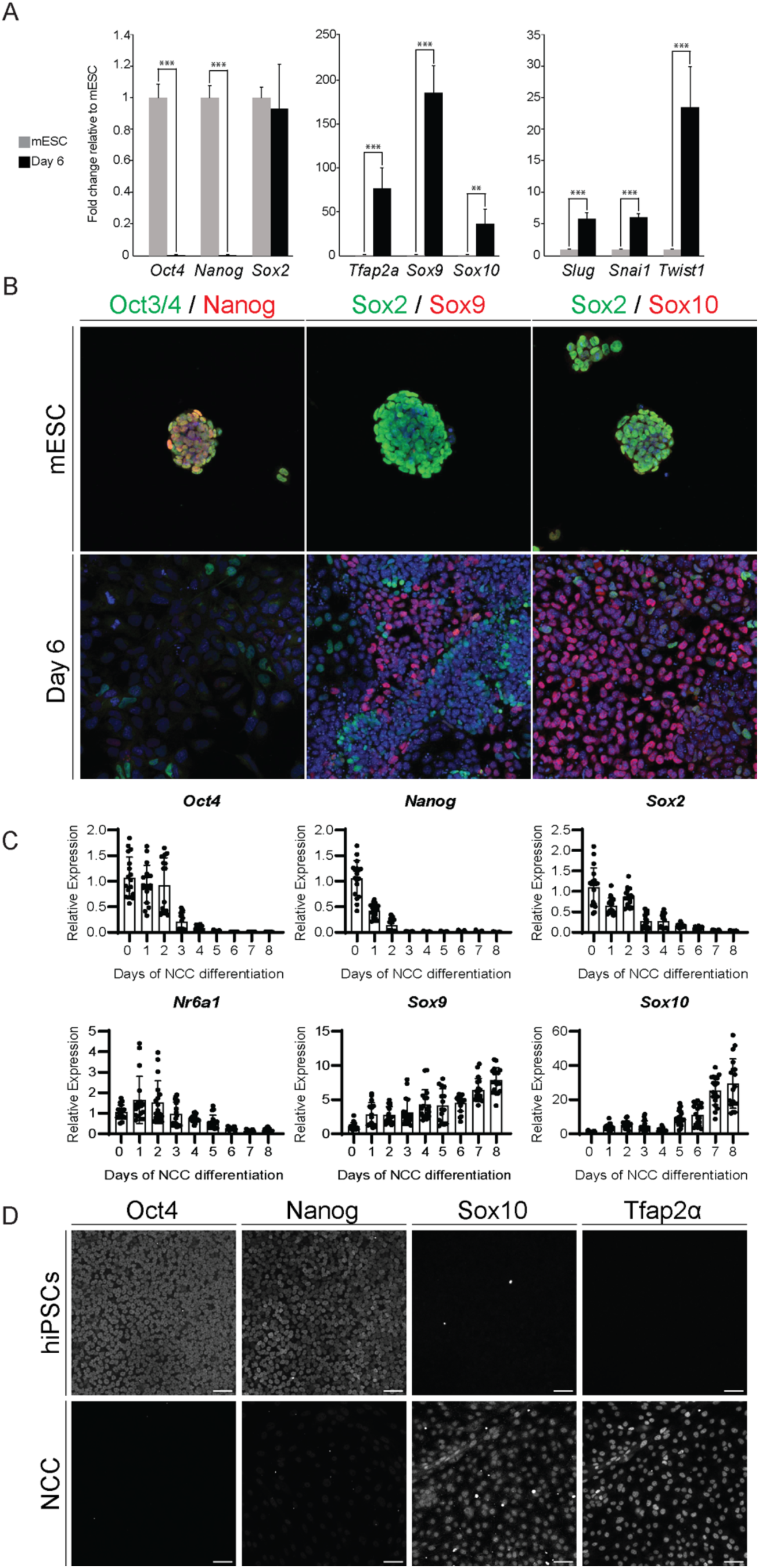
Pluripotency-associated factors are immediately downregulated upon inducing the differentiation of stem cells into NCC. A-B. mESCs were induced to become NCC through the provision of FGF to culture media. Samples were collected prior to induction (mESCs) and 6 days of differentiation, when cells are differentiated and considered NC-like. **A.** RT-qPCR analyses of samples at mESC and Day6 stages for pluripotency (*Oct4, Nanog* and *Sox2)*, NCC specification (*Tfap2a*, *Sox9* and *Sox10*) and EMT (*Snai1*, *Slug and Twist1*). Expression levels are normalized to mESC controls (day 0). At day 6, cells exhibit a downregulation of *Oct4* and *Nanog* and an upregulation of NCC-specifier and EMT genes. In contrast, mESCs do not express the NCC-specifier and EMT factors. **B.** Immunostaining of OCT3/4, NANOG, SOX2, SOX9 and SOX10 on mESCs and at 6 days of differentiation. mESCs express OCT4, NANOG and SOX2, but after 6 days of differentiation, the NC-like cells do not express these factors. **C-D.** hiPSCs induced to NCC in a 8-day differentiation culture. **C.** RT-qPCR analyses of hiPSCs for pluripotency (*Oct4*, *Nanog* and *Sox2*), NCC differentiation (*Sox9* and*Sox10*), and *Nr6a1*. Expression levels are normalized to hiPSC controls (day 0). *Oct4* and *Nanog* are downregulated upon the provision of differentiation media and expression is lost by Day 3. *Nr6a1* expression is highest during days 1-2, consistent with a function as a repressor of pluripotency and stem cell maintenance and with activating NCC specification and EMT. **D.** Immunostaining of OCT3/4, NANOG, SOX10 and TFAP2A in hiPSCs and at day 8 of NCC differentiation (NCC). OCT4 and NANOG are no longer expressed in the day 8 cells where NCC factors are expressed.

